# Spatially clustered piRNA genes promote the transcription of piRNAs via condensate formation of the H3K27me3 reader UAD-2

**DOI:** 10.1101/2023.12.10.571043

**Authors:** Chengming Zhu, Xiaoyue Si, Xinhao Hou, Panpan Xu, Jianing Gao, Yao Tang, Chenchun Weng, Mingjing Xu, Qi Yan, Qile Jin, Jiewei Cheng, Ke Ruan, Ying Zhou, Ge Shan, Demin Xu, Xiangyang Chen, Shengqi Xiang, Xinya Huang, Xuezhu Feng, Shouhong Guang

**Author notes:** These authors contributed equally to this work (Chengming Zhu, Xiaoyue Si).

## Abstract

PIWI-interacting RNAs (piRNAs) are essential for maintaining genome integrity and fertility in various organisms. In flies and nematodes, piRNA genes are encoded in heterochromatinized genomic clusters. The molecular mechanisms of piRNA transcription remain intriguing. Through unique molecular indexed-small RNA sequencing and chromosome editing, we discovered that spatial aggregation of piRNA genes enhances their transcription in nematodes. The heterochromatinized piRNA genome recruits the piRNA transcription complex USTC (including PRDE-1, SNPC-4, TOFU-4, and TOFU-5) and the H3K27me3 reader UAD-2, which phase separate into droplets to initiate piRNA transcription. We searched for factors that regulate piRNA condensate formation and isolated the SUMO E3 ligase GEI-17 as inhibiting and the SUMO protease TOFU-3 as promoting condensate formation, thereby regulating piRNA production. Our study revealed that spatial aggregation of piRNA genes, phase separation and deSUMOylation may benefit the organization of functional biomolecular condensates to direct piRNA transcription in the heterochromatinized genome.

## Introduction

PIWI-interacting RNAs (piRNAs) belong to a class of small noncoding RNAs that form complexes with PIWI proteins to repress transposable elements and regulate gene expression ^1–9^. In *Caenorhabditis elegans*, piRNAs originate from two large genomic clusters located on chromosome Ⅳ ^10–12^. These piRNA clusters are enriched with the repressive histone mark H3K27me3 ^13^. piRNAs are independently transcribed as short capped transcripts by RNA polymerase Ⅱ, generating piRNA precursor transcripts ^10–12^. The <u>u</u>pstream <u>s</u>equence transcription <u>c</u>omplex (USTC), which is essential for piRNA transcription, binds to the upstream Ruby motif of type Ⅰ piRNAs and promotes piRNA precursor transcription ^14^. The USTC complex consists of four proteins: PRDE-1, SNPC-4, TOFU-4 and TOFU-5 ^10, 14, 15^. These proteins generate distinct subnuclear foci that colocalize with piRNA clusters in the nucleus ^14^. The chromodomain protein UAD- 2 recognizes the histone mark H3K27me3 and facilitates the recruitment of the USTC complex to piRNA genes ^16–18^. However, despite our knowledge of piRNA clusters and heterochromatic states being crucial for piRNA transcription, the mechanisms remain mysterious ^13, 16, 19–23^. The underlying mechanism of USTC assembly at piRNA loci and the activation of transcription in repressive heterochromatic regions are yet to be fully understood.

In *Drosophila* ovarian germ cells, piRNAs are produced from dual-strand clusters embedded in heterochromatin marked by H3K9me3. The Rhino-Deadlock-Cutoff (RDC) complex binds to heterochromatin and licenses noncanonical transcription of dual-strand piRNA clusters. Rhino’s chromodomain recognizes H3K9me3 ^24–26^. *Drosophila* ovarian somatic cells also possess uni-strand piRNA clusters that depend on the Nxf1-Nxt1 complex for nuclear export of spliced piRNA precursors ^22, 27^. However, it remains unclear how piRNA genes evolved into specialized genomic clusters and how heterochromatin readers such as Rhino and UAD-2 interface with the core transcriptional machinery ^16, 26^.

In eukaryotes, biomolecular condensates form via liquid‒liquid phase separation and play crucial roles in various processes ^28–30^. Phase separation results from multivalent weak interactions between disordered regions in proteins and nucleic acids and leads to the formation of membraneless organelles ^31–34^. Recent studies have revealed that the phase separation of transcription factors can mediate the formation of transcriptional hubs ^35–37^, and condensation of activators and cofactors into phase- separated compartments potentiates gene expression ^38–41^. Hence, understanding whether and how phase separation underpins the compartmentalization of the heterochromatinized piRNA genome and transcriptional machinery will provide vital insights into the mechanism of gene regulation.

Small ubiquitin-like modifier (SUMO) is an essential posttranslational modification that regulates diverse cellular processes, including transcription, DNA repair, RNA processing, ribosome biogenesis, cell cycle control, and nuclear body formation ^42, 43^. Moreover, SUMOylation orchestrates the activity of transcription factors and phase separation of chromatin modifiers to control gene expression programs ^44–46^. SUMO-dependent phase separation facilitates the assembly of distinct subnuclear structures such as promyelocytic leukemia (PML) nuclear bodies ^38, 44^. SUMOylation of piRNA pathway proteins provides a molecular switch to connect piRNA target recognition to downstream chromatin silencing machinery, enabling heterochromatin formation and transcriptional silencing of piRNA target genes ^47–51^. Defining the mechanisms of SUMOylation-mediated control of piRNA transcription condensate formation will provide critical insights into the molecular underpinnings of piRNA production and genome regulation.

In this work, we show that the spatial clustering of piRNA genes into dense genomic loci promotes piRNA transcription in *C. elegans* through phase separation of the piRNA regulator UAD-2 into condensates at piRNA clusters. The regulated SUMOylation serves as a critical determinant directing UAD-2 condensation and piRNA synthesis. Our findings present a mechanistic model wherein piRNA gene clustering density, UAD-2 phase separation properties, and SUMOylation status collaboratively assemble functional piRNA transcription compartments.

## Results

### Clustered piRNA genes promote piRNA expression in *C. elegans* and *C. briggsae*

The *C. elegans* genome contains approximately 15,000 annotated piRNA genes distributed across six chromosomes, with ∼92% located within the two major piRNA clusters on chromosome Ⅳ (Extended Data Fig. 1a) ^11^. piRNA Cluster Ⅰ is a 2.5 Mb region on the center of chromosome Ⅳ, containing ∼200 piRNA genes per 100 kb. piRNA Cluster Ⅱ is a 3.7 Mb region on the right arm of chromosome Ⅳ, containing ∼400 piRNA genes per 100 kb (Fig. 1a) ^13, 52^. We use the SYP-1 protein of the synaptonemal complex as a marker to label the chromosomes in pachytene cells of the gonad ^53^. Through fluorescence imaging, we observed that the H3K27me3 reader UAD- 2 and the piRNA transcription factor TOFU-4 form a single nuclear focus concentrated on chromosome Ⅳ, corresponding to the Cluster II genome, in the nucleus of each pachytene germ cell (Fig. 1b,c) ^14, 16^. In contrast, Cluster Ⅰ and out-cluster regions exhibited minimal, if detectable, GFP foci. Chromatin immunoprecipitation (ChIP) revealed enriched binding of UAD-2 and the USTC components (TOFU-4 and TOFU- 5) on the Cluster Ⅱ genome compared to Cluster Ⅰ and out-cluster regions (Fig. 1d).

**Fig. 1.**
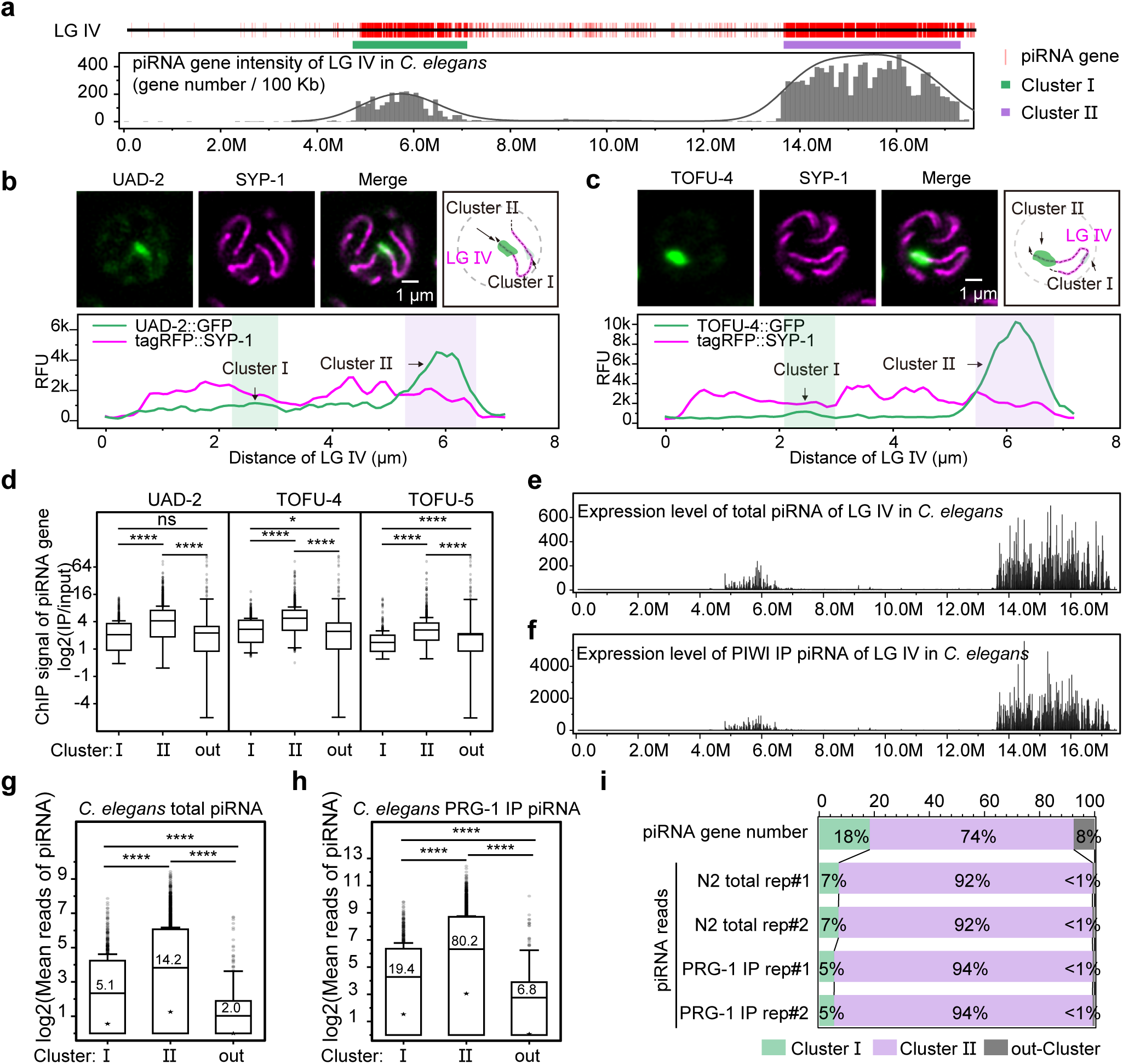
Clustered piRNA genes promote piRNA expression in *C. elegans*. **a,** The upper section displaying the localization of piRNA genes (marked in red) on *C. elegans* chromosome Ⅳ. The green and purple segments indicate Cluster Ⅰ and Cluster Ⅱ loci, respectively. The lower panel illustrates the piRNA gene density on chromosome Ⅳ, representing the number of piRNA genes per 100 kb segment. **b,** The upper section displaying the localization of UAD-2::GFP and tagRFP::SYP-1. The lower section illustrating the relative fluorescence unit intensity (RFU) of UAD-2 and SYP-1 along chromosome Ⅳ, marked by the dashed line. **c,** The upper section displaying the localization of TOFU-4::GFP and tagRFP::SYP-1. The lower section illustrates the relative fluorescence unit intensity (RFU) of TOFU-4 and SYP-1 along chromosome Ⅳ, marked by the dashed line. **d,** Boxplots revealing the ChIP signal (log_2_(IP/input)) of piRNA genes for the indicated protein. Statistical significance was determined using a two-sample t test. **e,** Graphs presenting the expression levels of total piRNAs from LG IV, based on average reads from two replicates, normalized as reads per million+1. **f,** Graphs presenting the expression levels of PRG-1-associated piRNAs from LG IV, based on average reads from two replicates, normalized as reads per million+1. **g,** Boxplots revealing log_2_(average piRNA reads of two replicates) for total piRNA across Cluster Ⅰ, Cluster Ⅱ, and out-Cluster regions. Normalized as reads per million+1. Statistical significance was determined using a two-sample t test. **h,** Boxplots revealing log_2_(average piRNA reads of two replicates) for PRG-1-associated piRNAs across Cluster Ⅰ, Cluster Ⅱ, and out-Cluster regions. Statistical significance was determined using a two-sample t test. Normalized as reads per million+1. **i,** Graphs presenting the fractional distribution of piRNA genes and their corresponding expression reads (normalized as reads per million) across Cluster Ⅰ, Cluster Ⅱ, and out- Cluster loci.

To characterize piRNA expression, we developed a unique molecular index (UMI)-based small RNA sequencing method and analyzed the piRNA populations from total RNA or PRG-1-associated RNA in wild-type adults (Extended Data Fig. 1b). The expression pattern of piRNAs across chromosome Ⅳ based on total small RNA and PRG-1-immunoprecipitated small RNA displayed enriched expression in the two clusters (Fig. 1e,f). piRNAs in Cluster Ⅱ showed substantially higher expression levels than those in Cluster Ⅰ and out-cluster regions. Specifically, Cluster Ⅱ piRNAs exhibited an average expression of approximately 14.2 reads per million (RPM) in the total RNA sequencing and ∼80.2 RPM in PRG-1-associated RNA sequencing. In contrast, Cluster Ⅰ piRNAs exhibited lower expression, while out-cluster piRNAs displayed further reduced expression (Fig. 1g,h). Among all piRNA genes, the Cluster Ⅰ and out-Cluster genomes contained 18% and 8%, respectively, of the total number of piRNA genes and expressed a minor fraction of piRNA reads (Fig. 1i). In comparison, Cluster Ⅱ contains 74% of piRNA genes and expresses close to 92% (total RNA) and 94% (PRG-1- associated RNA) of piRNA reads (Fig. 1i).

To further test whether the clustering of piRNA genes is important for piRNA expression, we analyzed the distribution and expression of piRNAs in another nematode, *C. briggsae*. The *C. briggsae* genome contains over 25,000 piRNAs across 6 chromosomes ^54^, clustering into one locus on LG Ⅰ and two loci on LG Ⅳ, which are termed Clusters 1, 2, and 3, respectively (Extended Data Fig. 1c). The published *C. briggsae* piRNA high-throughput sequencing data ^54^ showed that, similar to that of *C. elegans*, piRNA genes located within clusters exhibited explicitly higher expression compared to those of out-cluster regions. The average expression level of clustered piRNAs bound to the PIWI protein was ∼50 RPM, while out-cluster piRNAs showed an average of ∼3.1 RPM (Extended Data Fig. 1d). Further analysis revealed that 32% of piRNA genes were encoded in out-cluster regions while generating only 2% of the total piRNA reads. In contrast, 68% of the piRNA genes were clustered and contributed 98% of the total piRNA reads (Extended Data Fig. 1e).

Interestingly, a recent work has described a piRNA-mediated RNA interference system (piRNAi) operating via an extrachromosomal array in *C. elegans* ^55^, in which a short genomic region (1.5 kb) that contained at least six highly expressed piRNAs and were repeated approximately one hundred times in extrachromosomal arrays could promote efficient piRNA transcription. Notably, the piRNA density in these extrachromosomal arrays amounts to approximately 400 piRNAs per 100 kb, closely paralleling that observed in Cluster Ⅱ of *C. elegans*. Thus, these data suggested that the clustering of piRNA genes into dense genomic regions may be critical for robust piRNA expression in both *C. elegans* and *C. briggsae*.

### Intrachromosomal inversion alters piRNA expression

To test whether piRNA expression levels rely on gene density rather than their positions on chromosomes, we designed dual sgRNA-guided CRISPR/Cas9 technology to introduce chromosome inversion on LG IV ^56^. We designed 4 sgRNAs targeting the middle of Cluster Ⅰ and Cluster Ⅱ to cleave LG Ⅳ and screened for lines with intrachromosomal inversions in these regions using PCR (Fig. 2a). We obtained a strain in which Cluster Ⅰ-A was fused with Cluster Ⅱ-C and Cluster Ⅰ-B was fused with Cluster Ⅱ-D, therefore forming two new clusters, termed Cluster (A+C) and Cluster (B+D) (Fig. 2a and Extended Data Fig. 2a). In the new strain, the piRNA gene numbers and densities were similar between the two new clusters. Using UAD-2::GFP as the fluorescence reporter for piRNA genome location, we observed that the LG IV inversion strain exhibited two smaller foci on the same chromosome per nucleus compared to the presence of a single UAD-2 focus in each pachytene stage germ cell nucleus in wild-type worms (Fig. 2b). UMI-small RNA sequencing revealed that the proportion of piRNAs from the original Cluster Ⅰ increased from 4% in wild-type animals to 14% in the inverted strain, approaching the proportion of piRNA gene numbers (Fig. 2c and Extended Data Fig. 2b). Most piRNAs located in Cluster Ⅰ exhibited over 2-fold increased expression, with average expression rising from ∼2.9 RPM in wild-type to ∼8.7 RPM in inverted animals (Fig. 2d-2g, and Extended Data Fig. 2c,d). Furthermore, piRNAs located on the chromosomal arm or center locations (Cluster Ⅱ-C and Ⅱ-D) did not show significant expression changes (Extended Data Fig. 2e,f).

**Fig. 2.**
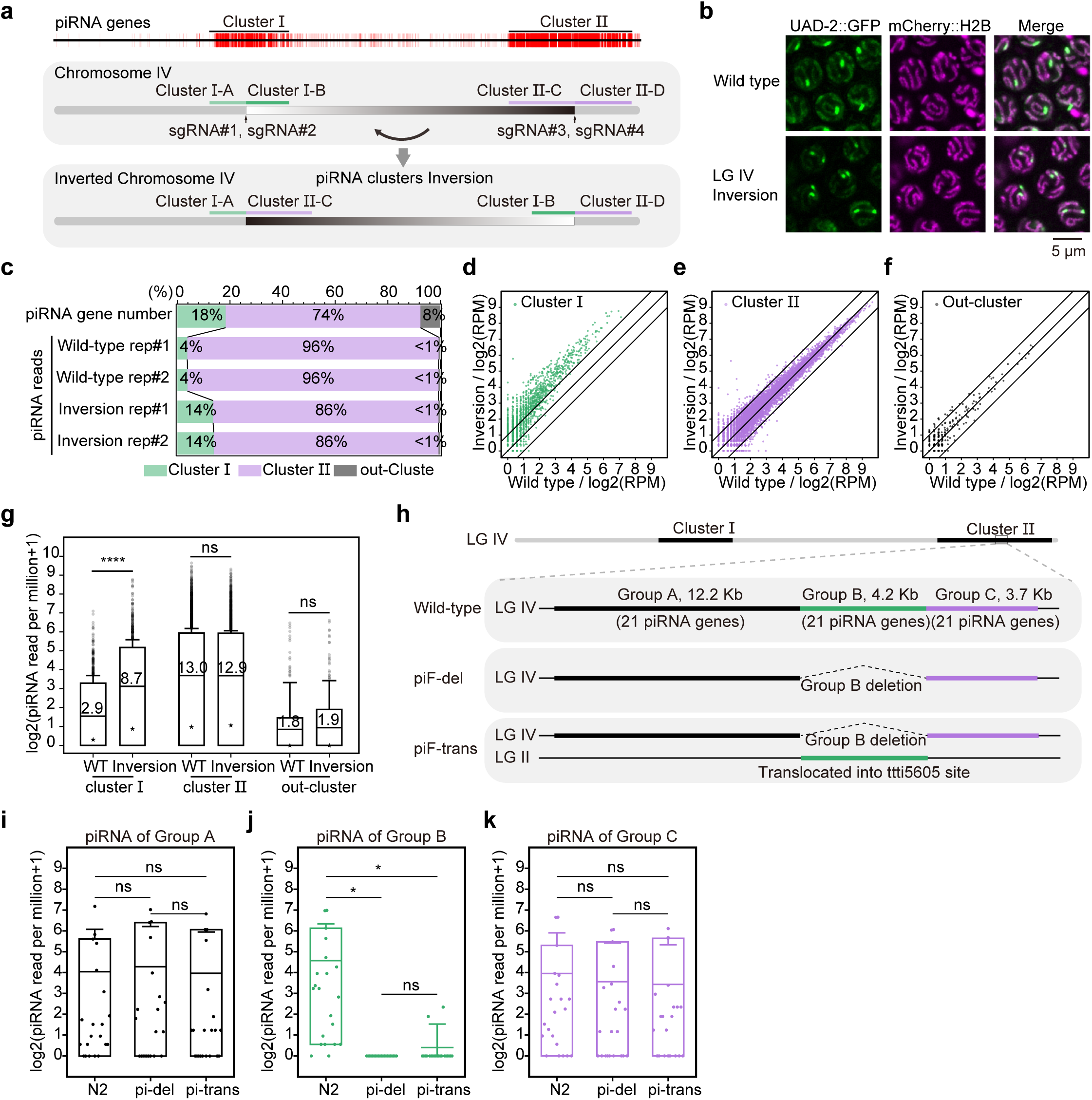
Chromosomal editing alters piRNA clustering and expression. a, Schematic diagram displaying the CRISPR-mediated intrachromosomal inversion process. sgRNA#1 and #2 target the center of Cluster Ⅰ, and sgRNA#3 and #4 target the center of Cluster Ⅱ. The lower panel illustrates the inverted LG Ⅳ chromosome of the worm. **b,** Images showing the subcellular localization of UAD-2::GFP and mCherry::H2B in the nuclei of pachytene germ cells. In wild-type cells, a single UAD- 2 bright spot is observed in each nucleus, whereas the inverted strain reveals two UAD- 2 bright spots per nucleus. **c,** Graphs presenting the fractional distribution of piRNA genes and their corresponding expression reads (normalized as reads per million) for the indicated strains across Cluster Ⅰ, Cluster Ⅱ, and out-Cluster loci. **d-f,** Scatter plots displaying the expression levels of Cluster Ⅰ, Cluster Ⅱ, and out-Cluster piRNAs in the wild-type (x-axis) and inversion strain (y-axis), based on the log_2_(average piRNA reads of two replicates), normalized as reads per million+1. **g,** Boxplot revealing log_2_(average piRNA reads of two replicates) for both wild-type and inversion strains across Cluster Ⅰ, Cluster Ⅱ, and out-Cluster loci. Normalized as reads per million+1. Statistical significance was determined using a paired-sample t test. **h,** Schematic diagram showing the CRISPR-mediated chromosomal translocation experiments. Group B piRNAs are translocated from Cluster Ⅱ of LG Ⅳ to the *ttti5605* site of LG Ⅱ. **i-k,** Boxplots revealing log_2_(normalized piRNA reads+1) for the wild-type, piRNA fragment deletion (piF-del) strain, and piRNA fragment translocation (piF-trans) strain across Group A, Group B, and Group C. Statistical significance was determined using a paired-sample t test.

### Interchromosomal translocation of the piRNA genome reduces piRNA expression

To test whether localization within a cluster is crucial for piRNA expression, we used CRISPR/Cas9 technology to delete a 4.2 kb segment (group B) from Cluster Ⅱ containing 21 actively expressed piRNA genes and inserted it into the ttti5605 site on chromosome Ⅱ (Fig. 2h). UMI piRNA sequencing showed that the Group B piRNAs exhibited nearly no expression in either the deletion (pi-del) or translocation (pi-trans) strains compared to wild-type animals. In contrast, the flanking segments, Group A and Group C, exhibited similar piRNA expression levels to those of wild-type animals (Fig. 2i-k).

Thus, these chromosome inversion and translocation experiments supported that the clustering of piRNA genes is crucial for robust piRNA transcription.

### Mobility properties of the H3K27me3 reader UAD-2 and the USTC complex

Fluorescence imaging revealed that UAD-2 and the USTC component TOFU-5 formed distinct subnuclear foci of approximately 1 μm^2^ within each germline nucleus at 20°C (Extended Data Fig. 3a,b). Our previous work showed that increasing temperature is detrimental to piRNA expression ^16^. When the worms were cultured at 25°C for approximately 24 hours, the piRNA transcriptional condensates dispersed. The piRNA foci recovered upon returning the worms to 20°C culture conditions (Extended Data Fig. 3c-f). We speculated that the clustered expression of piRNAs might be achieved through the condensation of certain transcription factors. Temperature might regulate piRNA transcription by modulating the aggregation and dispersion of piRNA transcription machinery.

**Fig. 3.**
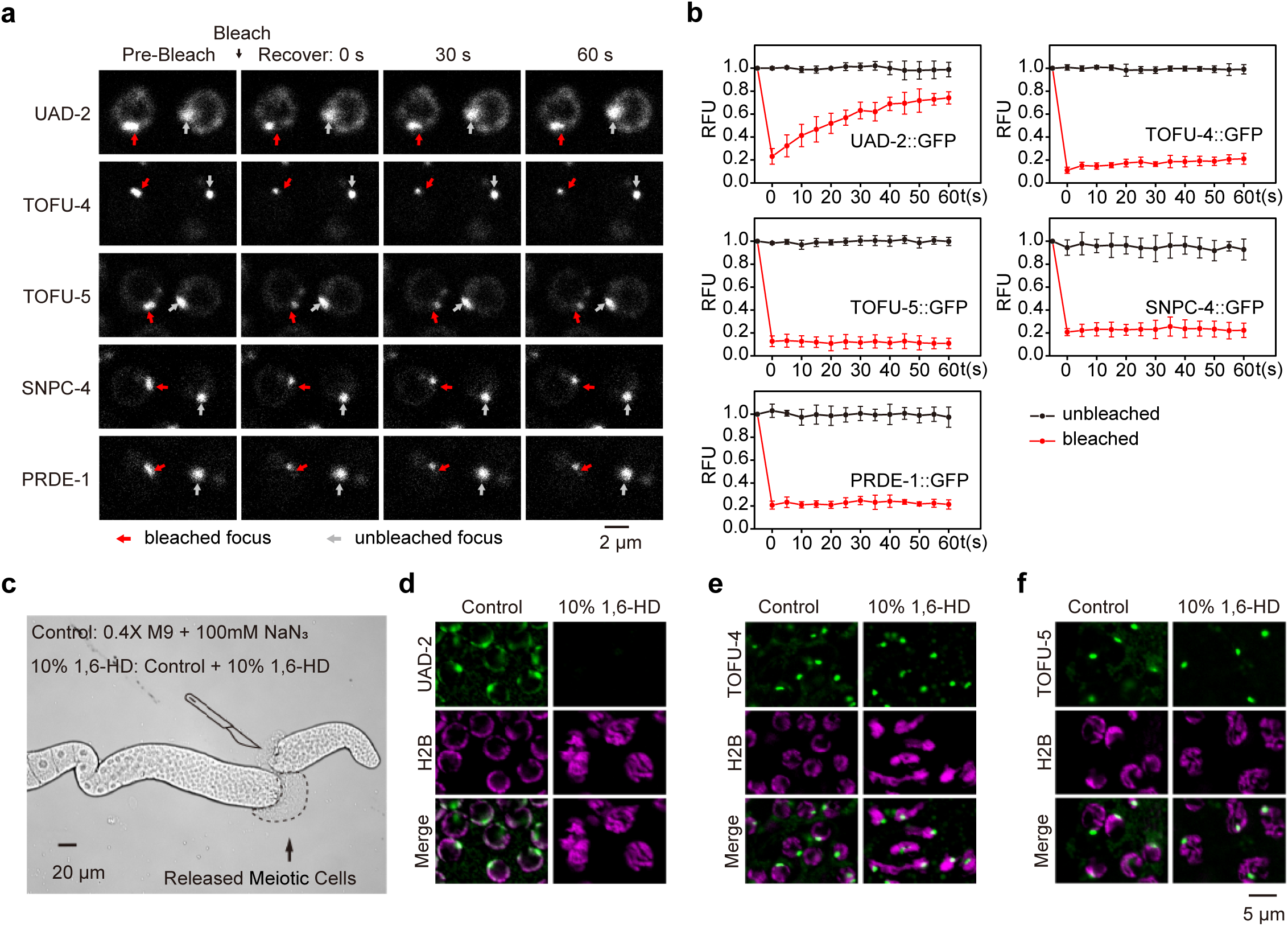
UAD-2 forms a liquid droplet-like condensate at piRNA cluster loci. a, Fluorescence recovery after photobleaching (FRAP) of UAD-2::GFP and GFP-tagged USTC components (TOFU-4, TOFU-5, SNPC-4 and PRDE-1) was conducted using a Zeiss LSM980 confocal microscope. **b,** Graphs presenting the relative fluorescence unit intensity (RFU) of the control area and bleached area of UAD-2 and the USTC components. n=6. **c-f,** Images showing germline nuclei expressing UAD- 2::GFP, TOFU-4::GFP or TOFU-5::GFP. After release by needle disruption, the germline was imaged within 5 minutes using a Leica Thunder imaging system. Germlines were treated with 10% 1,6-hexanediol to prohibit phase separation.

First, we assessed the protein mobility of piRNA transcription factors in vivo using fluorescence recovery after photobleaching (FRAP) assay. The rapid recovery of the photobleached UAD-2 aggregates revealed high mobility of UAD-2 in the nucleus (Fig. 3a,b). In contrast, the USTC components, including TOFU-4, TOFU-5, PRDE-1, and SNPC-4, displayed negligible, if any, recovery after photobleaching (Fig. 3a,b). Additionally, treating germ cell nuclei with 10% 1,6-hexanediol (1,6-HD) led to rapid dispersion of UAD-2 aggregates within 5 minutes, while the USTC foci were maintained (Fig. 3c-f). The protein mobility experiments suggested that the H3K27me3 reader protein UAD-2 might have a unique function in the dynamic regulation of piRNA transcriptional condensates.

### The intrinsically disordered regions may play key roles in the condensate formation of UAD-2

UAD-2 contains two intrinsically disordered regions (IDRs) and a chromodomain (Fig. 4a and Extended Data Fig. 4a). The chromodomain recognizes H3K27me3 modification. Intrinsically disordered regions are important to direct the formation of membraneless condensates through liquid-liquid phase separation (LLPS) ^9^. Using CRISPR/Cas9 technology, we deleted each of these domains and determined its subcellular localization and effect on piRNA expression. The deletion of the short and long IDRs partially disrupted UAD-2 foci, whereas the deletion of both IDRs nearly completely abolished focus formation and chromatin association (Fig. 4b). UAD- 2(ΔChromo) exhibited negligible, if any, expression, suggesting that the chromo domain is required for the stability of the UAD-2 protein (Fig. 4b).

**Fig. 4.**
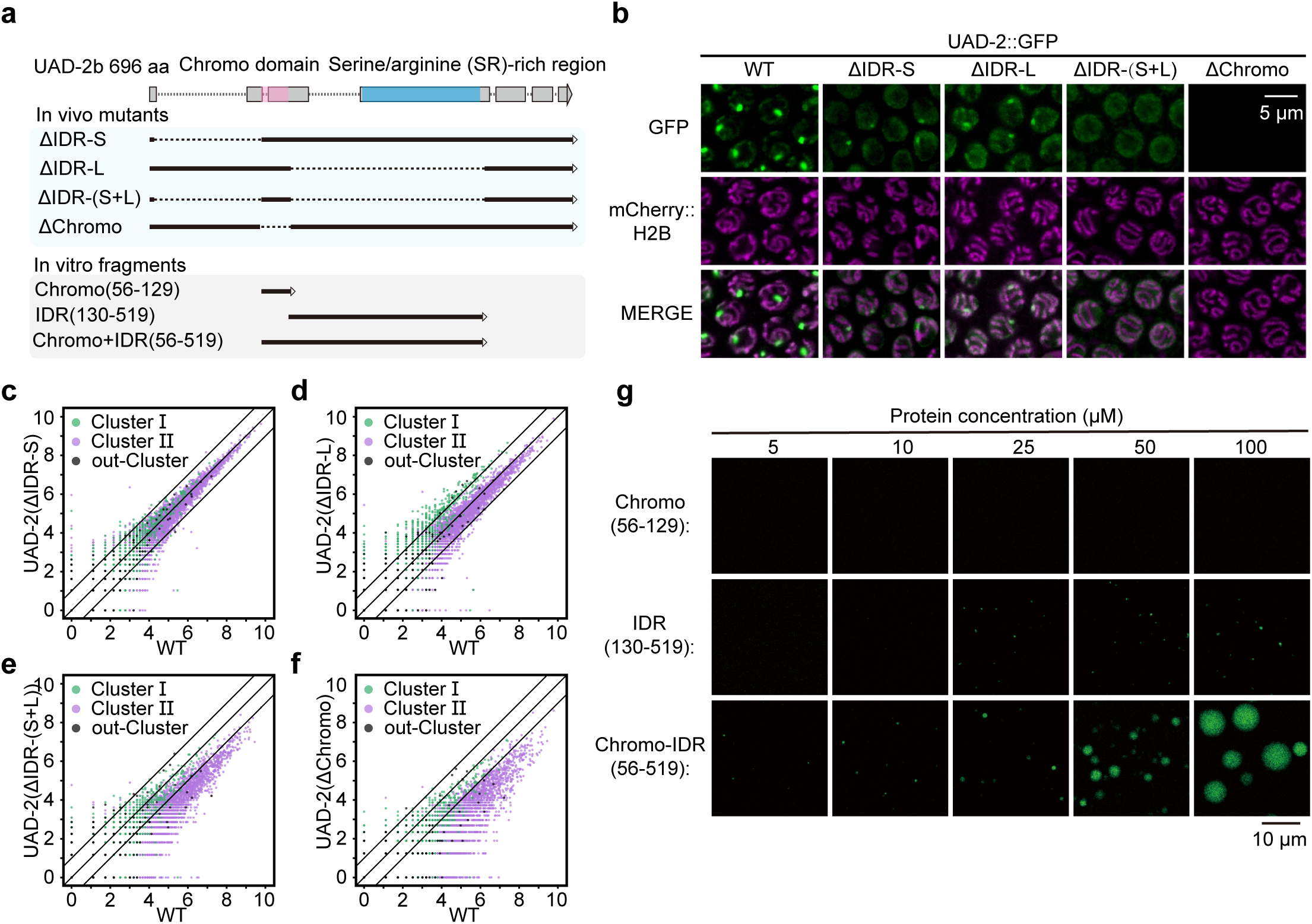
Intrinsically disordered regions of UAD-2 play a key role in condensate formation. a, Diagram illustrating the domain structure of UAD-2. The dash lines of ΔIDR-S, ΔIDR-L, ΔIDR-(S+L) and ΔChromo indicate short IDR sequence deletion, long IDR sequence deletion, double IDR sequence deletion and Chromo deletion in vivo, respectively. Chromo(56-129), Chromo+IDR(56-519) and IDR(130-519) indicate in vitro expressed fragments of UAD-2, respectively. **b,** Images showing the subcellular localization of the indicated UAD-2::GFP and mCherry::H2B transgenes in pachytene cells. **c-f,** Scatter plots displaying the expression levels of Cluster Ⅰ, Cluster Ⅱ, and out-Cluster piRNAs in the wild-type (x-axis) and indicated mutants (y-axis), based on the log_2_(reads per million+1). **g,** Images showing the liquid droplet formation of recombinant UAD-2 fragments in vitro.

In the *uad-2*(*ust200*) null allele animal, the USTC component PRDE-1 appeared largely dispersed within the nucleus (Extended Data Fig. 4b,c). The deletion of UAD-2 IDRs, unexpectedly, resulted in the appearance of an additional PRDE-1 focus, likely corresponding to the Cluster Ⅰ piRNA genome on LG IV.

The deletion of either the short or long IDR alone did not induce noticeable changes in piRNA levels (Fig. 4c,d). However, the deletion of both IDRs caused a dramatic downregulation of piRNAs, particularly those derived from the high-density Cluster Ⅱ loci (Fig. 4e). Furthermore, in the UAD-2(ΔChromo) mutants, piRNAs from Clusters Ⅰ and Ⅱ both decreased (Fig. 4f). These data suggest that the absence of the IDR sequence in UAD-2 leads to its dispersion and therefore the prohibition of piRNA expression.

To further investigate the role of the IDR sequence in UAD-2 in liquid-liquid phase separation, we expressed recombinant UAD-2 segments in *E. coli*, purified the proteins, and conducted an in vitro phase separation assay. The UAD-2 Chromo domain and IDR alone cannot efficiently form liquid droplets, while mixing the Chromo domain with IDR recombinant proteins induced the formation of phase-separated droplets at a 10 µM concentration (Fig. 4g), suggesting a synthetic effect of these two domains.

Collectively, these data suggested that UAD-2 may account for the mobility and phase separation ability in the condensation of piRNA transcription machinery.

### Forward genetic screening identified that GEI-17 suppresses piRNA expression

To identify the regulators of piRNA transcription, we performed two forward genetic screens to search for factors that bilaterally modulate the formation of UAD-2 aggregates. In pachytene germ cells, UAD-2 forms a distinct nuclear focus at 20°C and dissipates after heat shock for 24 hours under 25°C culture conditions (Extended Data Fig. 3c). We first mutagenized UAD-2::GFP animals with ethyl methanesulfonate (EMS) and searched for UAD-2 focus-retaining mutants via clonal screening at 25°C under fluorescence microscopy. From two thousand haploid genomes, we isolated two *gei-17* alleles (Fig. 5a-c and Extended Data Fig. 5a,b). Since GEI-17 is a highly conserved SUMO E3 ligase, we speculated that SUMOylation may prohibit the formation of piRNA foci (Fig. 5d). Indeed, knockdown of the SUMOylation gene *smo- 1* recovered the formation of UAD-2, TOFU-4 and TOFU-5 condensates at 25°C (Fig. 5e). Consistently, GEI-17 and SMO-1 suppressed piRNA expression at 25°C. In *gei- 17* mutants or *smo-1* knockdown animals, piRNA levels increased (Fig. 5f-h, and Extended Data Fig. 5c,d). GFP-tagged GEI-17 primarily localized to germ cell nuclei and colocalized with chromosomes, suggesting its capability to modify potential chromatin proteins and regulate piRNA expression (Fig. 5i). GEI-17 was also localized in embryos and somatic tissues (Extended Data Fig. 5e,f). Interestingly, GEI-17 was present in the nucleolar vacuoles in oocyte cells (Fig. 5i) ^57^.

**Fig. 5.**
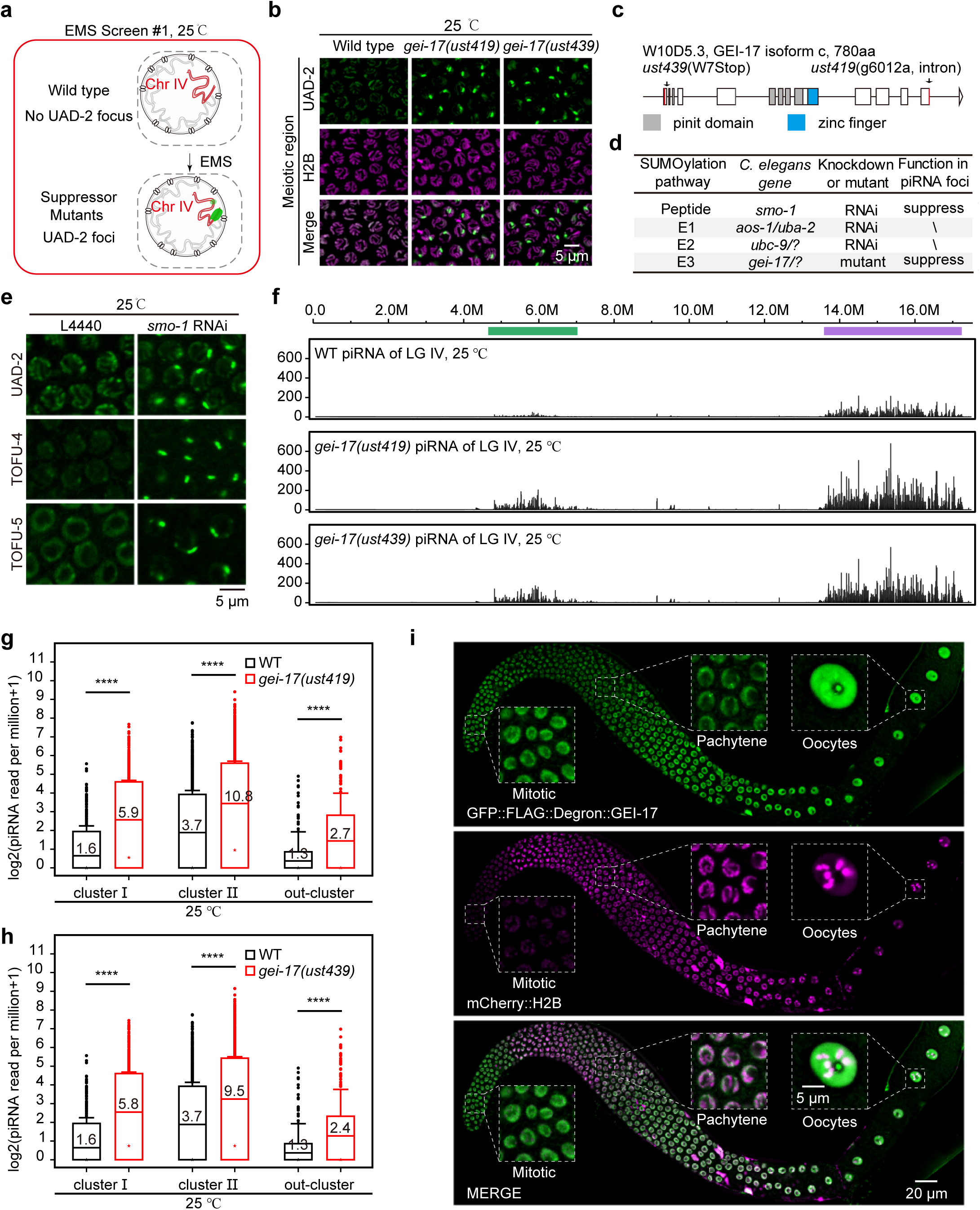
Forward genetic screening identifies GEI-17 suppressing piRNA expression. a, Schematic diagram displaying the forward genetic screening for UAD-2 focus suppressors in *C. elegans* germline nuclei at 25°C. **b,** Images showing pachytene cells of the indicated adult animals grown at 25°C. **c,** Diagram illustrating the *gei-17* exon and domain structure. The *ust439* allele changes tryptophan 7 to a stop codon and is likely a null allele. *ust419* modifies the 2nd base of the last intron. **d,** Summary of the SUMOylation pathway in *C. elegans*. **e,** Images showing pachytene cells of the dsRNA- treated animals grown at 25°C. **f,** Graphs presenting the expression levels of total piRNAs from LG IV of the indicated adult animals grown at 25°C, normalized as reads per million+1. **g and h,** Boxplots revealing log_2_(piRNA reads per million+1) of the indicated adult animals. Statistical significance was determined using a paired-sample t test. **i,** Images displaying the germline of adult animals. GFP::FLAG::Degron::GEI- 17 (green) partially colocalized with the chromatin marker mCherry::H2B (magenta).

Thus, these data suggested that SUMOylation suppresses the aggregation of piRNA transcription machinery and piRNA production at 25°C.

### The deSUMOylation peptidase TOFU-3 is required for the formation of UAD-2 condensates and piRNA production

We conducted a second forward genetic screening to search for factors that are required for UAD-2 condensation at 20°C. We mutagenized GFP::UAD-2 animals by EMS and isolated two *tofu-3* alleles, in which UAD-2 foci were dispersed in germ cells, via fluorescence microscopy-based clonal screening (Fig. 6a-c, and Extended Data Fig. 5a, 6a). A complementation assay between multiple *tofu-3* alleles and a transgene rescue experiment confirmed that TOFU-3 is essential for the formation of UAD-2 condensates (Extended Data Fig. 6b,c). *tofu-3,* also known as <u>u</u>biquitin-like <u>p</u>rotease (*ulp-5),* encodes a conserved deSUMOylation peptidase located on LG Ⅰ (Fig. 6c,d and Extended Data Fig. 6d). Knockdown of *ulp* family members showed that TOFU-3/ULP-5 is likely the only deSUMOylation peptidase involved in UAD-2 condensate formation (Extended Data Fig. 6e). Small RNA sequencing indicated that TOFU-3 is specifically required for the expression of the two clusters but not the out-cluster-localized piRNA genes (Fig. 6e,f and Extended Data Fig. 6f,g).

**Fig. 6.**
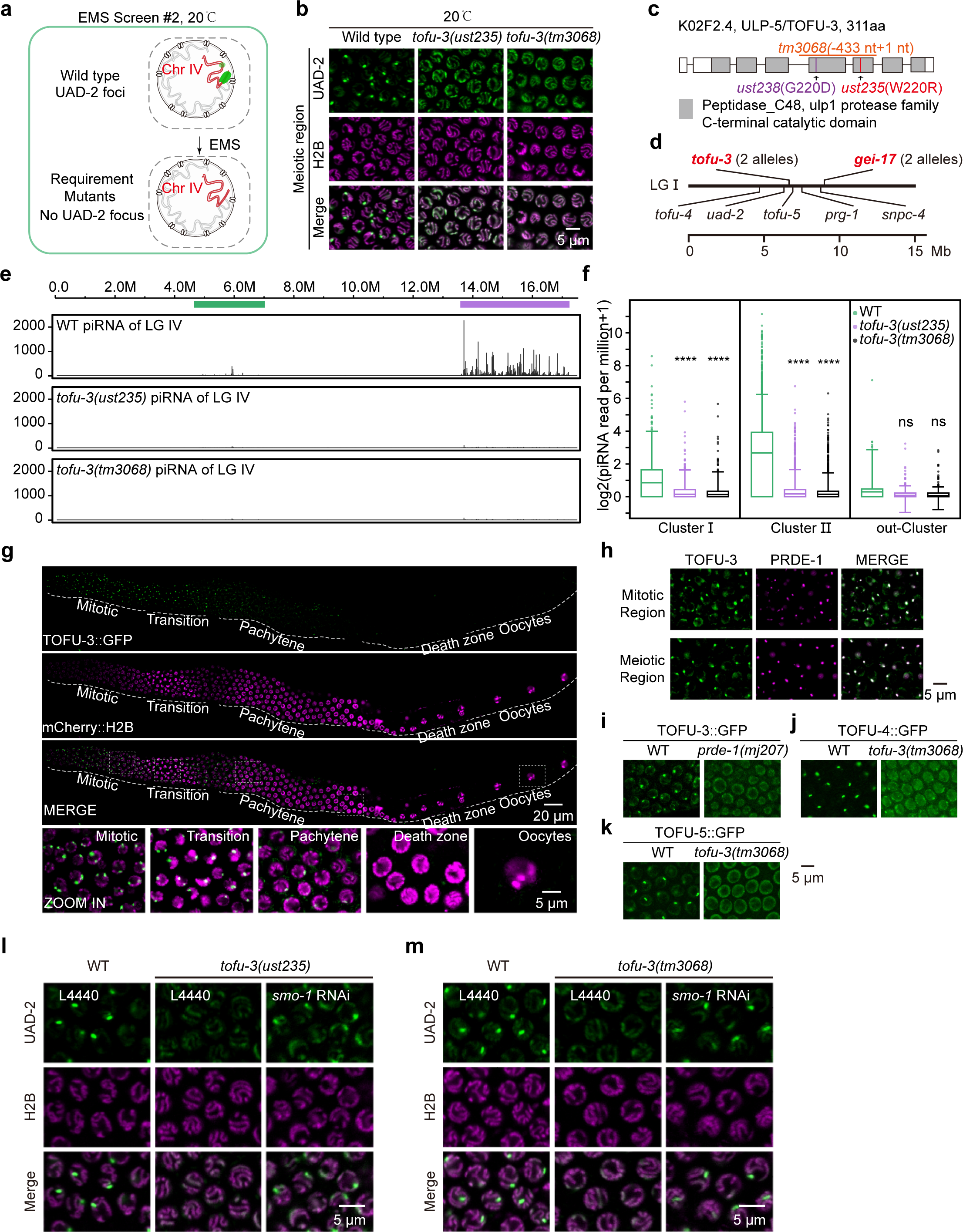
TOFU-3 is required for UAD-2 condensate formation and piRNA production. a, Schematic diagram displaying the forward genetic screening for mutants without UAD-2 focus in *C. elegans* germline nuclei at 20°C. **b,** Images showing pachytene cells of the indicated adult animals grown at 20°C. **c,** Diagram illustrating the *ulp-5/tofu-3* exon and domain structure. The alleles *ust238*, *ust235* and *tm3068* are highlighted. *tm3068* was likely a null allele and was used as a reference allele. **d,** Schematic representation summarizing piRNA genes and alleles identified from the two forward genetic screenings. **e,** Graphs presenting the expression levels of total piRNAs from LG IV of the indicated adult animals grown at 20°C, normalized as reads per million+1. **f,** Boxplots revealing log_2_(piRNA reads per million+1) of the indicated adult animals. Statistical significance was determined using a paired-sample t test. **g,** Images displaying the germline of adult animals. TOFU-3::GFP::3xFLAG (green) partially colocalized with the chromatin marker mCherry::H2B (magenta). **h,** Images showing colocalization of TOFU-3::GFP::3xFLAG (green) with mCherry::PRDE-1 (magenta) in mitotic and meiotic germline nuclei of adult animals. **i,** Images of representative meiotic germline nuclei of the indicated adult animals. TOFU-3::GFP::3xFLAG failed to form piRNA foci in *prde-1(mj207)* animals. **j and k,** Images of representative meiotic germline nuclei of the indicated adult animals. TOFU-4::GFP and TOFU- 5::GFP failed to form piRNA foci in *tofu-3(tm3068)* animals. **l and m,** Subcellular localization of UAD-2::GFP and mCherry::H2B in pachytene cells from WT worms fed L4440 and *tofu-3* mutant worms fed L4440 and *smo-1* dsRNA bacteria.

We generated an in situ 3xFLAG::GFP-tagged TOFU-3 by CRISPR/Cas9 technology. TOFU-3 was predominantly expressed in mitotic and transition zones, weakly expressed in early pachytene cells, and formed nuclear foci that colocalized with PRDE-1 (Fig. 6g,h). The mutation of *prde-1* prevents the formation of TOFU-3 foci (Fig. 6i). On the other hand, the formation of TOFU-4 and TOFU-5 foci also depends on TOFU-3 (Fig. 6j,k). These results suggested that TOFU-3 and the USTC complex exhibit mutual interdependency for localization, establishing TOFU-3 as an integral component of the piRNA transcription machinery.

To further test whether TOFU-3-mediated deSUMOylation is important for piRNA transcription, we knocked down *smo-1* via RNAi in *tofu-3* mutants. Remarkably, the depletion of *smo-1* rescued UAD-2 focus formation and substantially elevated piRNA expression in the *tofu-3* mutant background (Fig. 6l,m and Extended Data Fig. 6h).

We analyzed the fluorescence intensity profiles of UAD-2 and histone H2B. In wild-type nuclei, UAD-2 was prominently enriched at piRNA gene clusters but weakly colocalized with other chromatin regions. In *tofu-3* mutants, UAD-2 aggregation was ablated; instead, UAD-2 displayed substantially increased global chromatin association throughout the genome compared to that of wild-type animals (Extended Data Fig. 6i- k). These data suggested that the loss of TOFU-3 does not affect the chromatin binding ability of UAD-2 but likely disrupts its condensation ability.

### SUMOylation coordinates piRNA expression

At 20°C, wild-type germ cells displayed one explicit UAD-2 focus, corresponding to Cluster Ⅱ piRNA loci, in pachytene nuclei. However, we did not observe, if noticeable, a second explicit UAD-2 focus corresponding to Cluster Ⅰ piRNA loci (Fig. 1b and 7a). Strikingly, *gei-17* mutants displayed an additional pronounced UAD-2 focus corresponding to Cluster Ⅰ piRNA loci (Fig. 7b,c). Consistently, Cluster Ⅰ piRNAs were substantially upregulated in *gei-17* mutants at 20°C (Fig. 7d-f). Similarly, *smo-1* knockdown led to the formation of two UAD-2 foci and USTC foci in each pachytene nucleus (Extended Data Fig. 7a-f) and a significant upregulation of Cluster Ⅰ piRNA levels (Extended Data Fig. 7g,h).

**Fig. 7.**
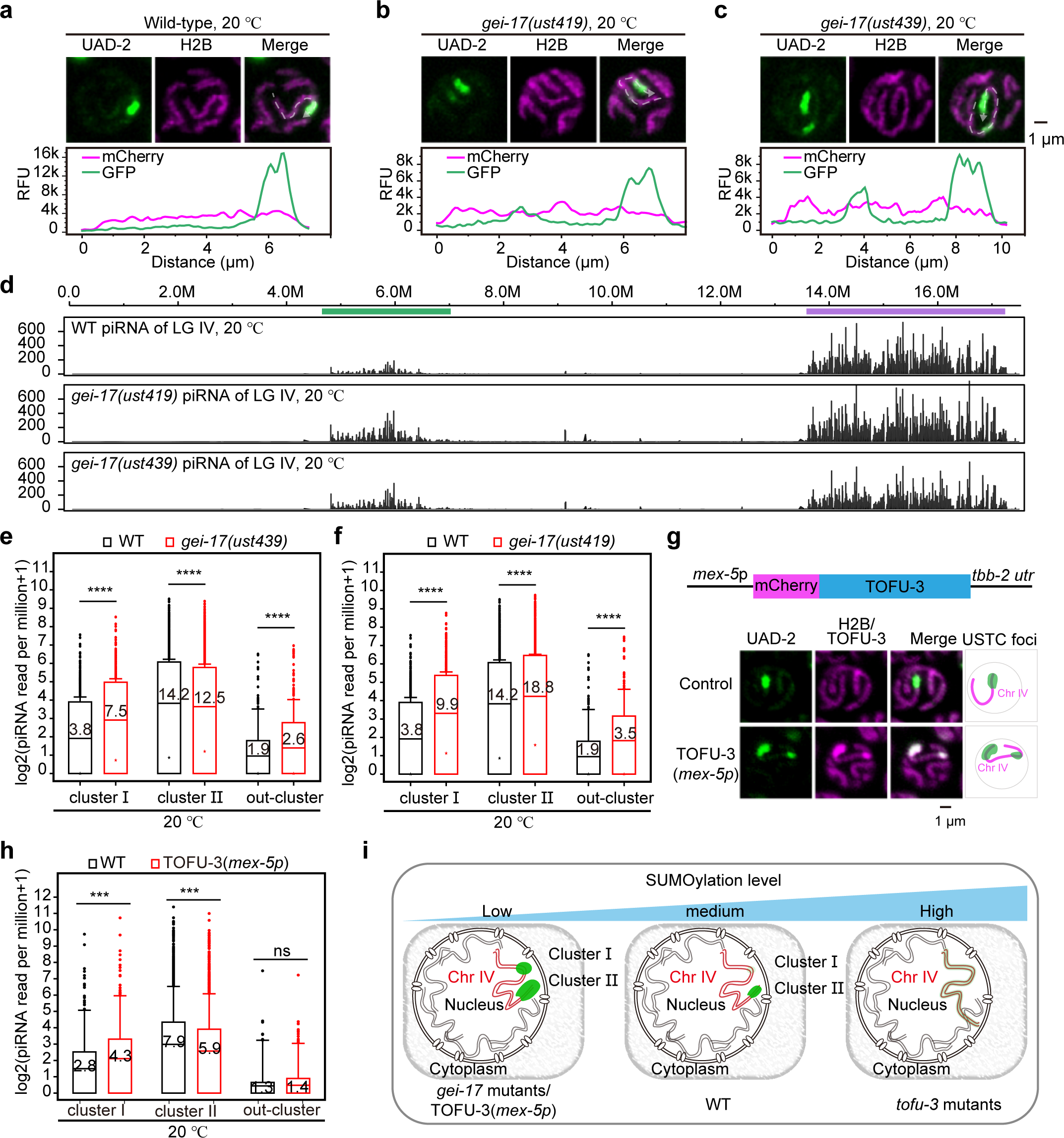
SUMOylation suppresses Cluster Ⅰ piRNA expression. a-c, (upper) Subcellular localization of UAD-2::GFP with mCherry::H2B in pachytene cells of the indicated animals maintained at 20°C. (lower) Relative fluorescence unit intensity (RFU) as indicated by dashed lines along chromosome Ⅳ in the upper panel. **d,** Expression levels of total piRNAs from LG IV of the indicated adult animals maintained at 20°C, normalized as reads per million+1. **e and f,** Boxplots presenting log_2_(piRNA reads per million+1) of the indicated adult animals. Statistical significance was determined using a paired-sample t test. **g,** (upper) Diagram illustrating mCherry::TOFU-3 by the *mex-5* promoter and *tbb-2* utr (TOFU-3(*mex-5p*)). (lower) Subcellular localization of UAD-2::GFP with mCherry::H2B/mCherry::TOFU-3 in pachytene cells of the indicated animals maintained at 20°C. **h,** Boxplots presenting log_2_(piRNA reads per million+1) of the indicated adult animals. Statistical significance was determined using a paired-sample t test. **i,** Schematic diagram of the SUMOylation level regulating UAD-2::GFP and USTC complex condensation.

To further investigate the role of SUMOylation in regulating piRNA expression across piRNA clusters, we introduced a single-copy construct with *mex-5* promoter-driven mCherry::TOFU-3 into the WT worm germline. The addition of mCherry::TOFU-3(*mex-5p*) in pachytene cells resulted in reduced SUMOylation at piRNA loci and enhanced UAD-2 binding at low-density piRNA regions when cultured at 20°C (Fig. 7g). Our piRNA sequencing data revealed a significant upregulation of piRNA expression from Cluster Ⅰ genes upon the addition of mCherry::TOFU-3(*mex- 5p*) (Fig. 7h). These findings indicate that SUMOylation selectively represses piRNA gene expression in regions with low piRNA gene density under standard culture conditions in WT animals.

Together, these data suggested that GEI-17-mediated SUMOylation and TOFU- 3-mediated deSUMOylation bilaterally modulate the condensation of piRNA transcription machineries and coordinate piRNA expression (Fig. 7i).

## Discussion

In this study, we showed that the density of piRNA gene clustering in the genome is critical to promote piRNA expression in *C. elegans*, via quantification of piRNA expression across nematode species and chromosome manipulation assays. Furthermore, we revealed that the piRNA factor UAD-2 undergoes liquid-liquid phase separation to form dynamic condensates enriched at piRNA gene clusters. In addition, SUMOylation plays pivotal roles in directing the assembly and modulating the dynamics of UAD-2 condensates. The SUMO protease TOFU-3 is indispensable for stimulating UAD-2 phase separation and piRNA production. In contrast, the SUMO E3 ligase GEI-17 antagonizes piRNA condensate integrity. Thus, regulated SUMOylation is likely a key molecular switch orchestrating UAD-2 condensation and piRNA transcription (Fig. 7i). In summary, we showed that piRNA gene density, UAD-2 phase separation properties, and SUMOylation status constitute an intricate regulatory circuit that cooperatively orchestrates piRNA transcription factory and piRNA production. We proposed that UAD-2 condensates at dense piRNA clusters and enhances transcription, a process fine-tuned by SUMOylation (Fig. 8a).

**Fig. 8.**
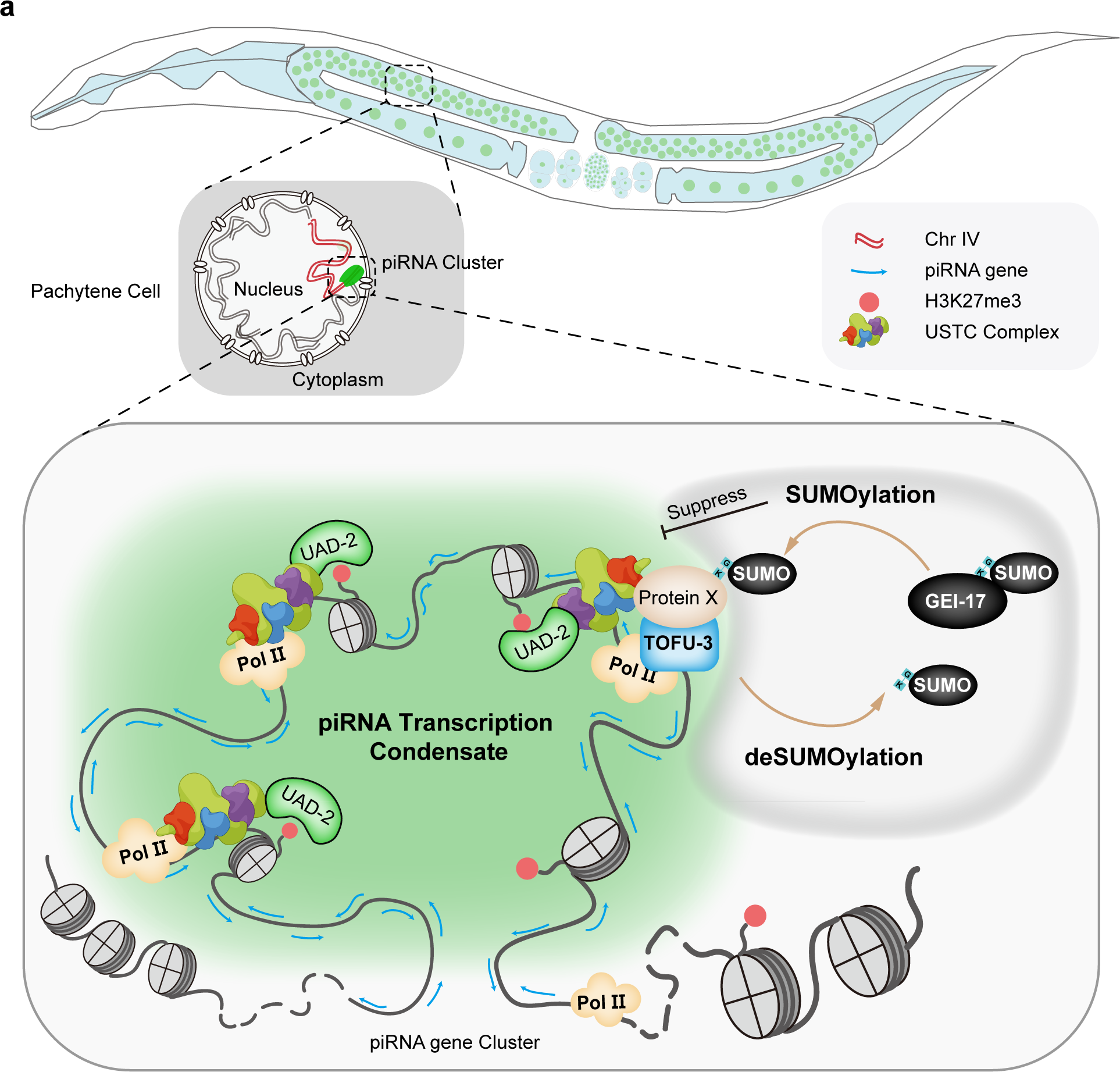
A working model for piRNA transcription in *C. elegans*. a, In *C. elegans*, densely clustered piRNA genes facilitate the assembly of UAD-2 condensates, which in turn actively recruit the USTC complex and POL Ⅱ and subsequently initiate piRNA transcription. Concurrently, the SUMO E3 ligase GEI-17 antagonizes piRNA condensate integrity, and TOFU-3-mediated deSUMOylation promotes piRNA condensate formation and piRNA biogenesis.

### piRNA transcription from the heterochromatinized genome

The mechanism of transcription from heterochromatic regions remains poorly understood. In *Drosophila* ovarian germ cells, piRNAs are produced from dual-strand clusters embedded in heterochromatin marked by H3K9me3, which is recognized by the chromo protein Rhino. Here, we found a similar clustering mechanism to maintain efficient piRNA transcription from H3K27me3-enriched genomic loci, which are recognized by the chromo protein UAD-2, in *C. elegans*. The unique biophysical properties of UAD-2, including high intrinsic disorder and mobility, allow it to access and dynamically regulate clustered piRNA loci. Patel *et al*. also reported from Hi-C data that piRNA clusters could form transcription hubs through long-distance interactions ^58^. Together, these results suggest that eukaryotes may use a universally conserved strategy to boost piRNA transcription via the spatial aggregation of piRNA genes to enhance gene density and proximity.

### Phase separation regulates transcriptional condensate formation

Transcription occurs in a compartmentalized manner via “transcription factories” to facilitate subnuclear localized transcription factors. Specific genomic clustering of short noncoding RNA genes may promote compartmentalization and efficient gene expression. Thus, piRNA transcription condensates provide an ideal model to dissect the mechanism(s) by which phase separation regulates transcription in vivo and how chromatin organization and posttranslational modifications, for example, SUMOylation, orchestrate gene expression. Integrative application of multidisciplinary approaches to address these questions will provide deeper mechanistic insights into the phase-separated control of noncoding RNA synthesis and transcription.

### SUMOylation and piRNA production

SUMOylation regulates liquid-liquid phase separation (LLPS) through multivalent interactions mediated by SUMO and SUMO interacting motifs (SIMs), promoting the formation of biomolecular condensates ^42, 46, 59^. In vitro experiments showed that combining polySUMO and polySIM results in phase-separated droplets. In vivo observations revealed that stress could induce SUMOylation and the assembly of biomolecular condensates, such as the promyelocytic leukemia (PML) nuclear body ^44, 60^. Our work revealed that in the germline cells of *C. elegans*, SUMOylation suppresses the formation of piRNA transcription condensates, thereby inhibiting the piRNA transcription process. We speculated that SUMOylation modifications on diverse protein substrates can induce a spectrum of physicochemical property shifts. Different SUMOylation types, such as poly-SUMO versus mono-SUMO, may confer distinct influences on biomolecular condensate formation. The intracellular environment is complex, and using *C. elegans* as a model system to study how SUMOylation regulates biomolecular condensates and transcription could provide significant value in exploring the in vivo biological functions of SUMOylation modification.

Previous studies in both *Drosophila* and *C. elegans* have identified the SUMOylation process as crucial for piRNA-mediated transcriptional silencing by suppressing target genes ^47–51^. However, our findings suggested that SUMOylation could also modulate piRNA production. While SUMOylation inhibits piRNA transcription, the specific deSUMOylation enzyme, TOFU-3, promotes piRNA transcription. Thus, SUMOylation may exert different regulatory roles across these separate processes. How SUMOylation serves as a repressive force in Pol II-driven piRNA expression processes requires further exploration.

piRNAs are non-coding small RNAs with specialized functions during distinct phases of germ cell development, necessitating meticulous transcriptional control. At developmental stages, it becomes imperative to precisely modulate piRNA transcription. In such scenarios, the transient binding affinity of UAD-2 combined with the suppressive effect of SUMOylation can effectively terminate piRNA transcription. Under specific environmental stresses, the inherent mobility of UAD-2 and the effects of SUMOylation may facilitate the organism’s adaptation through gene expression modulation.

### Limitations and perspectives of the study

While our study has advanced the understanding of piRNA regulation, several key questions remain. Identifying the substrate(s) of SUMOylation will illuminate the key molecular switches governing compartment dynamics. Elucidating the interaction and regulatory relationship between the histone reader UAD-2, the USTC complex, and the core transcription machinery is crucial for understanding the mechanism of piRNA

transcription. Additionally, investigating whether and how specific transcription factors are involved in piRNA transcription will greatly benefit the understanding of heterochromatic genome expression.

## Acknowledgments

We are grateful to the members of the Guang lab for their comments. We are grateful to the International *C. elegans* Gene Knockout Consortium and the National Bioresource Project for providing the strains. Some strains were provided by the CGC, which is funded by the NIH Office of Research Infrastructure Programs (P40 OD010440).

## Funding

This work was supported by grants from the National Key R&D Program of China (2022YFA1302700) and the National Natural Science Foundation of China (32230016, 32270583, 32070619, 2023M733425 and 32300438). This study was supported in part by the Fundamental Research Funds for the Central Universities.

## Author contributions

S.X., X. Huang, X.F. and S.G. conceptualized the research; C.Z., X. Huang, X.F. and S.G. designed the research; C.Z., X.S., P.X., Y.T., Q.J., K.R., Y.Z., G.S., D.X., X.C., and X. Huang performed the research; C.W., M.X., Q.Y. and J.C. contributed new reagents; C.Z., X. Hou and J.G. contributed analytic tools and performed bioinformatics analysis; C.Z., X.F. and S.G. wrote the paper.

## Competing interests

The authors declare no competing interests.

## Data Availability

The raw sequence data reported in this paper have been deposited in the Genome Sequence Archive in National Genomics Data Center, China National Center for Bioinformation / Beijing Institute of Genomics, Chinese Academy of Sciences (GSA: CRA012796) and are publicly accessible at https://ngdc.cncb.ac.cn/gsa.

## Extended Materials and Methods

### Strains

Bristol strain N2 was used as the standard wild-type strain. All strains were grown at 20°C unless otherwise specified. For the heat stress treatment, worms were cultured at 25°C. The strains used in this study are listed in Extended Data Table 1.

### Construction of deletion mutants and chromosome editing technology

For gene deletions, triple/quadruple-sgRNA–guided chromosome deletion was conducted as previously described ^1^. To construct sgRNA expression vectors, the 20 bp unc-119 sgRNA guide sequence in the pU6::unc-119 sgRNA(F+E) vector was replaced with different sgRNA guide sequences. Addgene plasmid #47549 was used to express the Cas9 Ⅱ protein. Plasmid mixtures containing 30 ng/µl of each of the three or four sgRNA expression vectors, 50 ng/µl Cas9 Ⅱ-expressing plasmid, and 5 ng/µl pCFJ90 were coinjected into UAD-2::GFP::3xFLAG (ustIS151) animals. Deletion mutants were screened by PCR amplification and confirmed by sequencing. The sgRNA sequences are listed in Extended Data Table 2.

Dual-sgRNA-mediated chromosome inversion and translocation were conducted as described previously ^2^.

### Construction of plasmids and transgenic strains

For the in situ transgene expressing GFP::3xFLAG tagged TOFU-4/SNPC- 4/PRDE-1/TOFU-3, a GFP::3xFLAG region was PCR-amplified from SHG326 genomic DNA. The coding sequence of GFP::3xFLAG was inserted before the stop codon using the CRISPR/Cas9 system. The sgRNA target sequences are listed in Extended Data Table 2. The ClonExpress MultiS One Step Cloning Kit (Vazyme C113- 02, Nanjing) was used to connect these fragments with the vector, which was amplified with 5′-TGTGAAATTGTTATCCGCTGG -3′ and 5′- CACACGTGCTGGCGTTACC-3′ from L4440. The injection mix contained PDD162 (50 ng/µl), repair plasmid (50 ng/µl), pCFJ90 (5 ng/µl), and two sgRNAs (30 ng/µl). The mix was injected into young adult N2 animals. The transgenes were integrated into the *C. elegans* chromosome by the CRISPR/Cas9 system.

### Forward genetic screening

Forward genetic screening experiments were conducted as previously described ^3^. Briefly, to identify the factors that can negatively or positively regulate UAD-2 focus formation, we mutagenized GFP::UAD-2 animals by ethyl methanesulfonate (EMS) and searched for mutants that either reformed UAD-2 foci at 25°C or depleted UAD-2 foci at 20°C by clonal screening. The F2 progeny worms were visualized under a fluorescence microscope at the gravid adult stage. Mutants revealing UAD-2 foci at 25°C or no UAD-2 foci at 20°C were selected. Both *gei-17* and *tofu-3* were identified by snp-SNP mapping followed by resequencing of the genome.

### RNAi

RNAi experiments were performed at 20°C or 25°C by placing synchronized embryos on feeding plates as previously described ^4^. HT115 bacteria expressing the empty vector L4440 (a gift from A. Fire) were used as controls. Bacterial clones expressing double-stranded RNAs (dsRNAs) were obtained from the Ahringer RNAi library and sequenced to verify their identity. All RNAi feeding experiments were performed for one generation from L1 to the gravid adult stage. Images were collected using a Leica DM4B microscope with a Leica Thunder image processing system.

### Fluorescence recovery after photobleaching (FRAP)

FRAP experiments were performed using a Zeiss LSM980 laser scanning confocal microscope at room temperature. Worms were anesthetized with 2 mM levamisole. A region of interest was bleached with 30% laser power for 5 ms, and the fluorescence intensities in these regions were collected every 5 s and normalized to the initial intensity before bleaching. For analysis, image intensity was measured by the mean and further analyzed by Origin software.

### 1,6-hexanediol treatment experiments

Adult worms were gently disrupted using a needle to release the germline nuclei. To preserve the integrity and activity of the released nuclei, imaging was promptly performed within a narrow window of 5 minutes post-disruption using a Leica DM4B microscope with a Leica Thunder image processing system. Dissected gonads were treated with 10% 1,6-hexanediol (referred to as 1,6-HD) supplemented with 0.4x M9 and 100 mM NaN3.

### Imaging

Images were collected using a Leica DM4B microscope with a Leica Thunder image process system. All worms were imaged at the gravid adult stage unless otherwise specified.

### Recombinant protein expression and purification

The UAD-2 chrome domain (amino acids 56-129), the UAD-2 IDR (amino acids 130-519) and the UAD-2 chrome domain+IDR (amino acids 56-129) were PCR-amplified and cloned and inserted into a plasmid (pET-28a-N8×H-MBP-3C vector) and expressed in *E. coli* BL21(DE3) cells. Protein expression was induced by 0.5 mM isopropyl β-d-1-thiogalactopyranoside (IPTG) at 16°C for 24 h. The recombinant

proteins were affinity purified through Ni-NTA His-binding resin (GE Healthcare), and the MBP tag was cleaved off using PreScission protease. The eluted proteins were further purified using size-exclusion chromatography with a SuperdexTM75 Increase 10/300 GL column or SuperdexTM200 Increase 10/300 GL column (GE Healthcare) on an ÄKTA pure chromatography system.

### In vitro UAD-2 droplet-formation assay

Purified recombinant proteins were incubated with the fluorochrome Alex488 (Thermo Fisher Scientific) overnight at 4°C and diluted to varying concentrations in buffer containing 100 mM NaCl, 25 mM NaH_2_PO_4_, 2 mM EDTA, and 5 mM DTT, pH 7.0. Protein solution (6 µl) was loaded onto a glass-bottom confocal dish (Biosharp). The droplet was imaged by a Zeiss LSM 980 confocal microscope with a 63× oil immersion lens.

### ChIP-Seq

ChIP-Seq experiments were conducted previously ^5, 6^, and the UAD-2, TOFU-4 and TOFU-5 ChIP-Seq data were downloaded from the China National Center for Bioinformation-National Genomics Data Center under submission numbers CRA004102 and CRA009179.

### Total RNA isolation

Synchronized late young adult worms were washed in M9 and ground with homogenizer lysis buffer (20 mM Tris⋅HCl [pH 7.5], 200 mM NaCl, 2.5 mM MgCl_2_, and 0.5% Nonidet P-40). The eluates were incubated with TRIzol reagent (Invitrogen) followed by isopropanol precipitation and DNase Ⅰ digestion (Qiagen).

### Isolation of PRG-1-associated RNA

Synchronized late adult GFP::3xFLAG::PRG-1 animals were ground by a homogenizer in lysis buffer (20 mM Tris-HCl [pH 7.5], 200 mM NaCl, 2.5 mM MgCl_2_, and 0.5% Nonidet P-40). The lysate was precleared with protein G-agarose beads (Roche) and then incubated with anti-FLAG M2 Magnetic Beads (Sigma #M8823). The beads were washed extensively, and GFP::3xFLAG::PRG-1 and associated RNAs were eluted with 100 μg/ml 3xFLAG peptide (Sigma). The eluates were incubated with TRIzol reagent (Invitrogen), followed by isopropanol precipitation.

### Unique molecular index (UMI)-small RNA sequencing

A total amount of 200 ng RNA per sample was used as input material for the small RNA library construction. The sequencing libraries were generated using a QIAseq miRNA Lib Kit following the manufacturer’s recommendations, and index codes were added to attribute sequences to each sample. Briefly, a preadenylated DNA adapter was ligated to the 3’ ends of miRNA, siRNA and piRNA. After the 3’ ligation reaction, another RNA adapter was then ligated to the 5’ end. The reverse transcription (RT) primer containing an integrated UMI subsequently binds to a region of the 3’ adapter and facilitates conversion of the 3’/5’ ligated miRNAs, siRNA and piRNA into cDNA while assigning a UMI to every small RNA molecule. During reverse transcription, a universal sequence is also added that is recognized by the sample indexing primers during library amplification. After reverse transcription, the cDNA was cleaned using a streamlined magnetic bead-based method. PCR amplification was performed using the universal forward and reverse primer-containing index. After library amplification, another cleanup of the library was performed using a streamlined magnetic bead-based method. Finally, library quality was assessed on the Agilent Bioanalyzer 2100 system using DNA High Sensitivity Chips. The clustering of the index-coded samples was performed on a cBot Cluster Generation System using TruSeq SR Cluster Kit v3-cBot-HS (Illumina) according to the manufacturer’s instructions. After cluster generation, the library preparations were sequenced on an Illumina platform, and 150 bp paired-end reads were generated. Raw data were transformed into clean reads by cutting the first 100 bases and removing low-quality reads. Subsequently, clean reads with correct UMI patterns were extracted using UMI-tools v1.0.0, serving as the basis for all downstream analyses to ensure high quality. Finally, the extracted reads were mapped to the reference sequence with Bowtie and deduplicated using UMI tools based on mapping coordinates and the attached UMI.

### Small RNA sequencing

Small RNAs were subjected to deep sequencing using an Illumina platform (Novogene Bioinformatics Technology Co., Ltd). Briefly, small RNAs ranging from 18 to 30 nt were gel-purified and ligated to a 3′ adaptor (5′- pUCGUAUGCCGUCUUCUGCUUGidT-3′; p, phosphate; idT, inverted deoxythymidine) and a 5′ adaptor (5′-GUUCAGAGUUCUACAGUCCGACGAUC-3′). The ligation products were gel-purified, reverse transcribed, and amplified using Illumina’s small RNA (sRNA) primer set (5′-CAAGCAGAAGACGGCATACGA-3′; 5′-AATGATACGGCGACCACCGA-3′). The samples were then sequenced using an Illumina Hiseq platform.

### RNA-seq analysis

The Illumina-generated raw reads were first filtered to remove adaptors, low-quality tags, and contaminants to obtain clean reads at Novogene. For piRNA analysis, clean reads ranging from 17 to 35 nt were mapped to the *C. elegans* transcriptome assembly WS243 using Bowtie2 with default parameters. The numbers of reads targeting each transcript were counted using custom Perl scripts. piRNAs were counted using custom Python scripts dependent on the piRNA annotation from Extended Data Table 3. The number of total reads mapped to the transcriptome minus the number of total reads corresponding to sense ribosomal RNA (rRNA) transcripts (5S, 5.8S, 18S, and 26S) was used as the normalization number to exclude the possible degradation fragments of sense rRNAs.

### piRNA gene annotations

piRNA annotations and genomic coordinates of piRNA genes were obtained by SAMtools against the *C. elegans* ce10 genome assembly. Information on piRNA clusters and piRNA Ruby-motif scores was obtained from a previous publication ^7, 8^.

### Statistics

Bar graphs with error bars are presented as the mean and SD. All of the experiments were conducted with independent *C. elegans* animals for the indicated N times. For the boxplots of ChIP-seq and piRNA expression level analysis, the box itself represented the interquartile range between 5% and 95%, thereby encompassing the middle 90% of the data set. The central horizontal line within the box and the adjacent numeric value within the box signify the arithmetic mean of the data. Outliers are individually marked as points, representing data points that fall outside the range of mean±1.5 standard deviations (SD). Statistical analysis is described in the following figure legends.

## Extended Data Figure legends

**Extended Data Table 1 List of strains used in the work.**

**Extended Data Table 2 Sequences of sgRNAs for CRISPR Cas9-mediated gene editing.**

**Extended Data Table 3 piRNA gene reference.**

**Extended Data Fig. 1.**
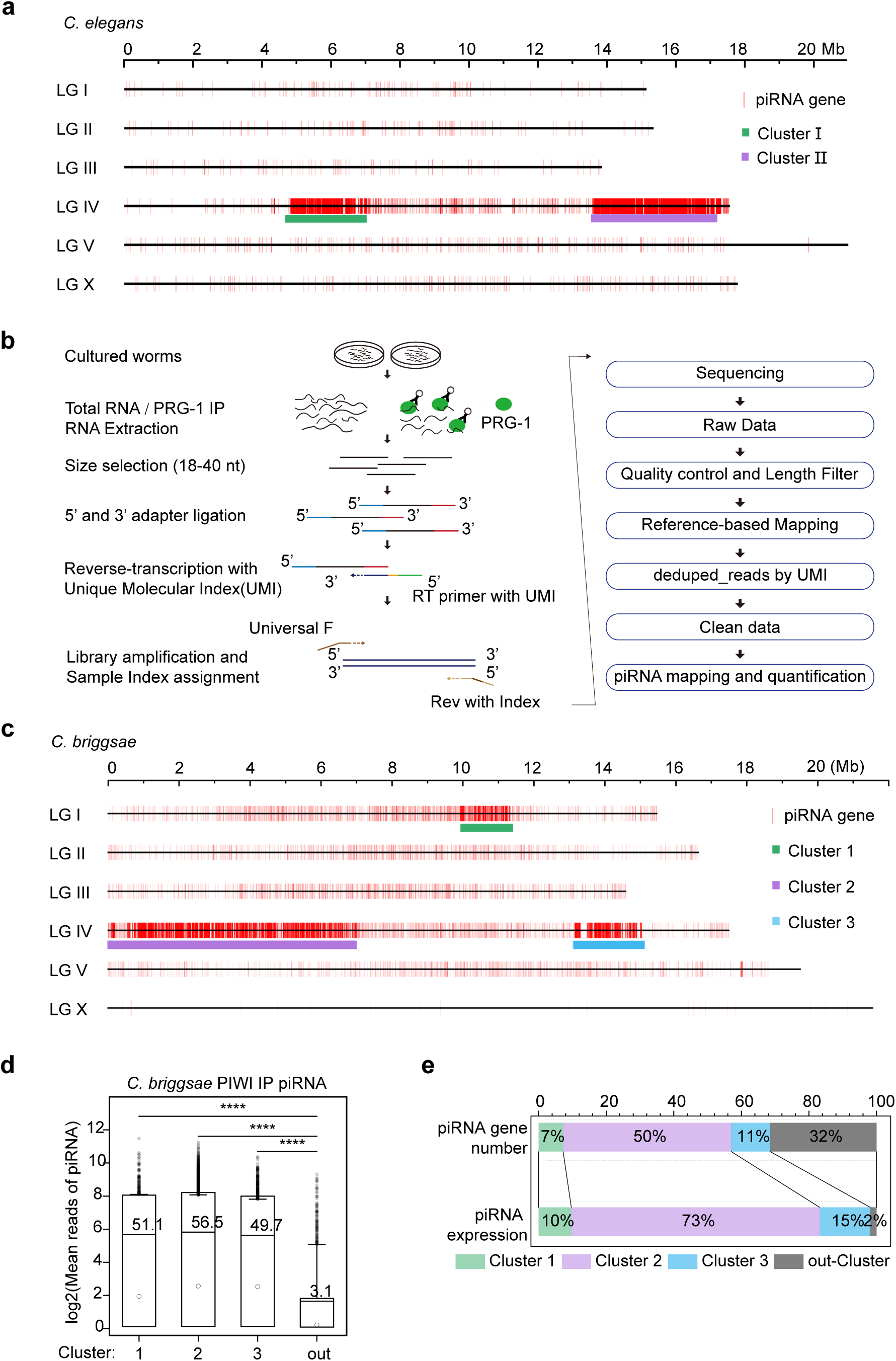
Spatial clustering of piRNA genes promotes piRNA expression independent of the Ruby motif. a, Genome-wide distribution of piRNA genes (n=15365) in *C. elegans*. **b,** A flowchart for UMI piRNA sequencing. **c,** Genome-wide distribution of piRNA genes (n=25883) in *C. briggsae*. **d,** Boxplots presenting log_2_(mean of piRNA reads from two replicates) for Cbr-PRG-1-associated piRNAs across Cluster 1, Cluster 2, Cluster 3, and out-Cluster in *C. briggsae*. Statistical significance was determined using a two-sample t test. **e,** Graphs presenting the fractional distribution of piRNA genes and their corresponding expression reads (normalized as reads per million from two replicates) across Cluster 1, Cluster 2, Cluster 3, and out-Cluster in *C. briggsae*.

**Extended Data Fig. 2.**
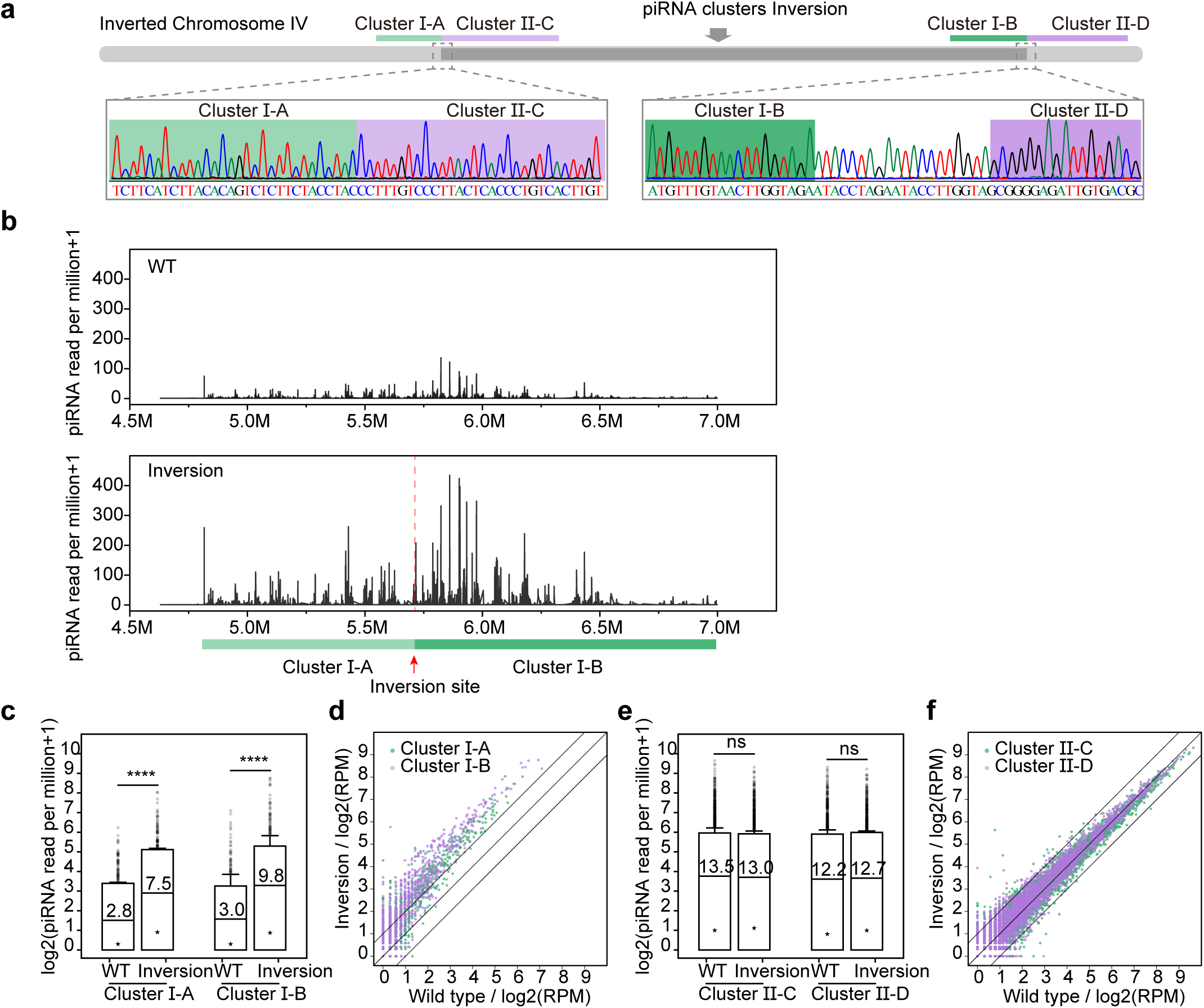
CRISPR/Cas9-mediated chromosome inversion and translocation alter piRNA expression. a, Diagram showing sequence details of intrachromosomal inversion in LG IV. **b,** Graphs presenting the expression levels of total piRNAs from Cluster Ⅰ of the indicated animals, based on average reads from two replicates, normalized as reads per million+1. **c-f.** Expression levels of piRNAs from various clusters between wild-type and inverted strains, presented using both boxplots and scatter plots. All expression values are the average of two replicates and are normalized as reads per million+1. Statistical significance was determined using a paired-sample t test.

**Extended Data Fig. 3.**
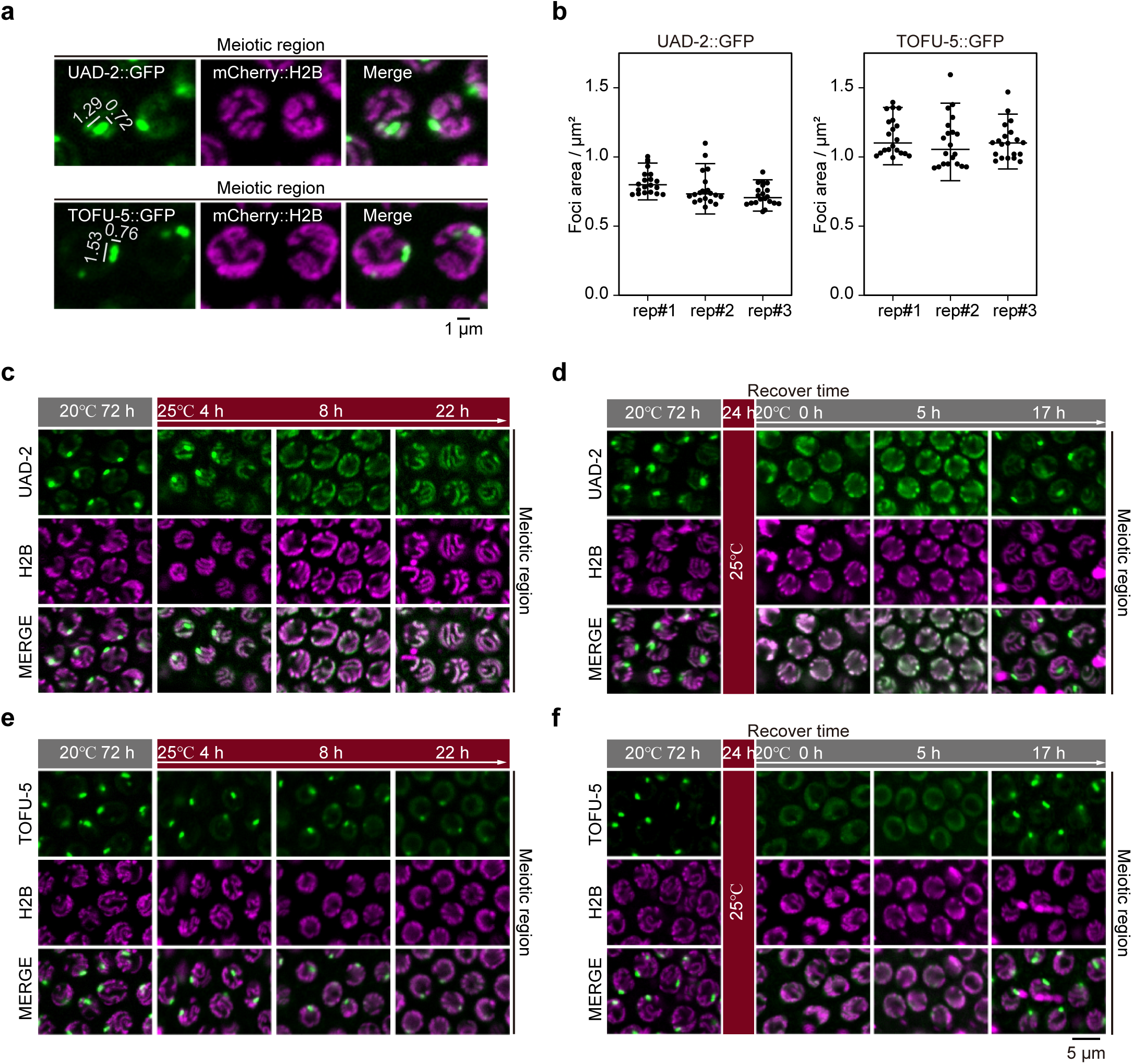
UAD-2 and the USTC complex form liquid droplet-like condensates at piRNA cluster loci. a, The upper panel shows the localization of UAD-2::GFP and mCherry::H2B in pachytene cells. The lower panel shows the localization of TOFU-5::GFP and mCherry::H2B in pachytene cells. **b,** Statistical analysis of the focus area for UAD- 2::GFP and TOFU-5::GFP in pachytene cells. n≥15 for each worm. **c-f,** The images reflect the subcellular localization of the mentioned proteins in pachytene cells under the indicated culture conditions.

**Extended Data Fig. 4.**
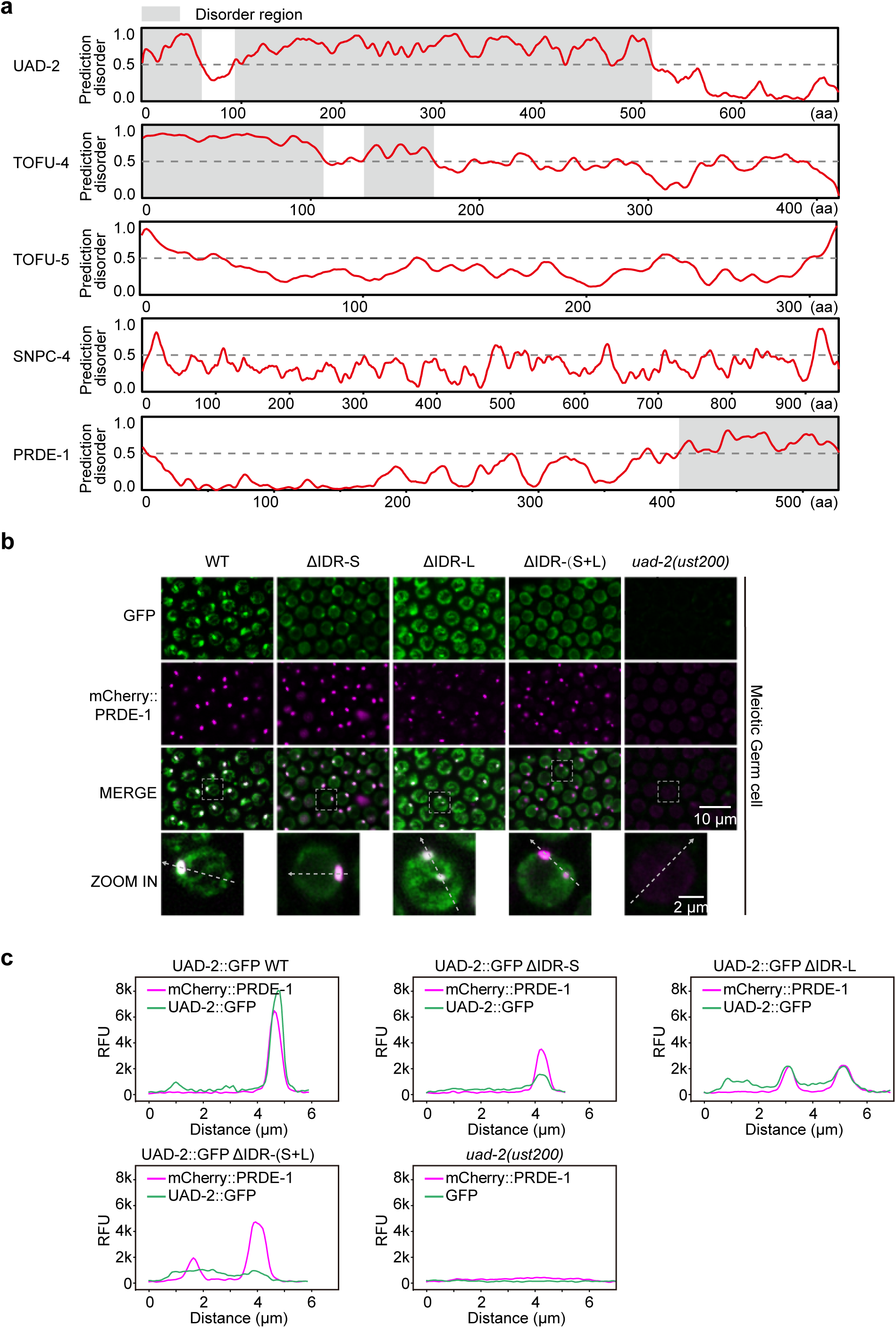
Phase separation property of UAD-2. a, IUPred2 prediction of UAD-2 and the USTC components using the IUPred2 long disorder option. **b,** Images showing the subcellular localization of the indicated UAD- 2::GFP variants in pachytene cells. **c,** Graphs presenting the relative fluorescence unit intensity (RFU) of the indicated UAD-2 variants, marked by the dashed line.

**Extended Data Fig. 5.**
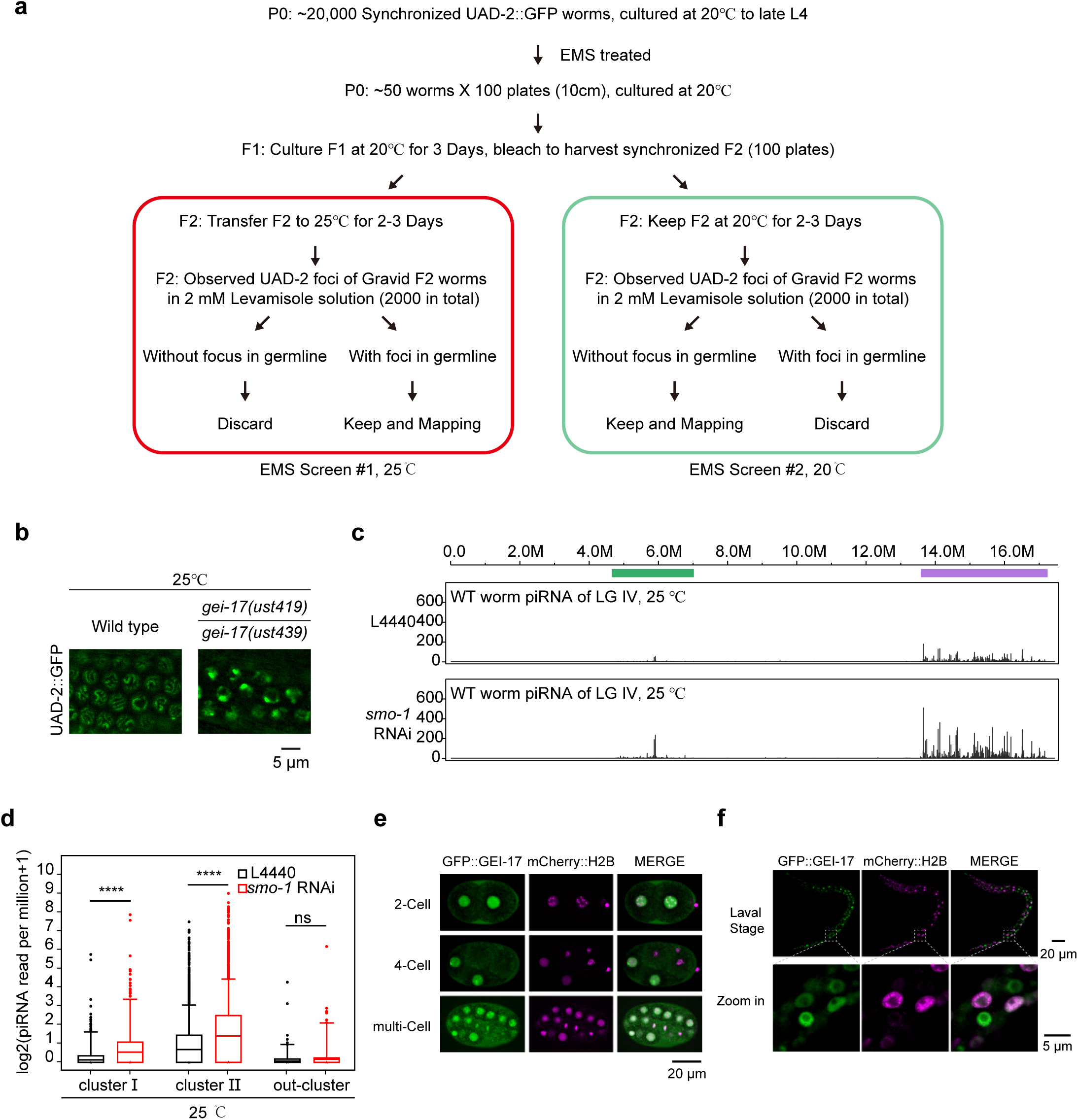
Forward genetic screening identifies factors suppressing UAD-2 foci formation. a, Forward genetic screening procedures for UAD-2::GFP regulators at 20°C and 25°C. **b,** Complementation test for two alleles of *gei-17* at 25°C. **c,** Expression levels of total piRNAs from LG IV in WT worms fed L4440 and *smo-1* dsRNA at 25°C, normalized as reads per million+1. **d,** Boxplots illustrating log_2_(piRNA reads per million+1) for WT worms fed with L4440 and *smo-1* dsRNA at 25°C. Statistical significance was determined using a paired-sample t test. **e and f,** Images displaying GFP::FLAG::Degron::GEI-17 (green) and mCherry::H2B (magenta) in embryos and larval animals.

**Extended Data Fig. 6.**
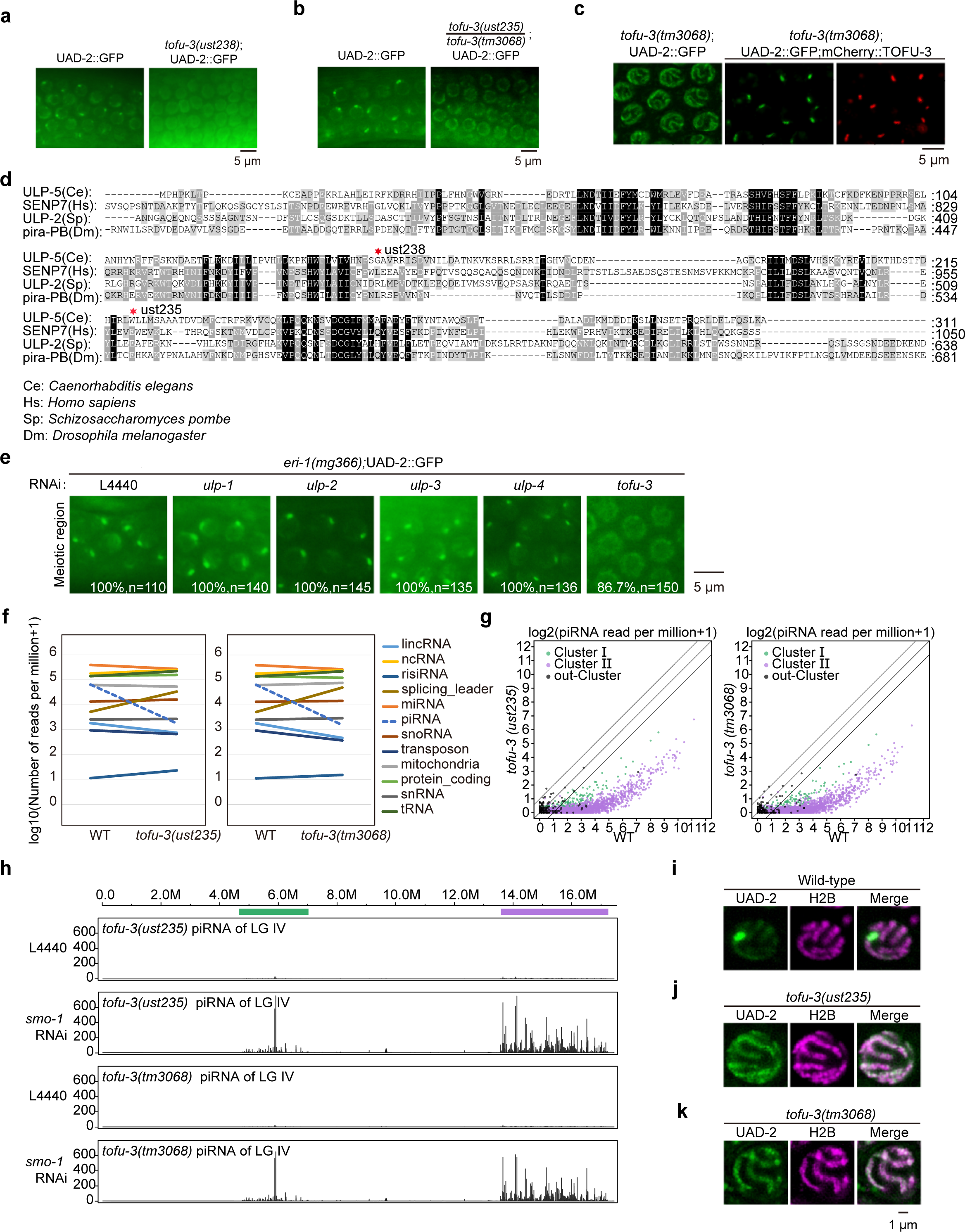
Forward genetic screening identified TOFU-3 as required for UAD-2 condensate formation and piRNA production. a. Images of the meiotic region of the indicated adult animal germline grown at 20°C. **b,** Complementation test of two alleles of *tofu-3* at 20°C. **c,** Images of pachytene cells of the indicated adult animals grown at 20°C. mCherry::TOFU-3 rescued UAD-2::GFP foci formation in *tofu-3(tm3068)* cells. **d,** Sequence alignment of the TOFU-3 (ULP-5) protein. **e,** Subcellular localization of UAD-2::GFP in pachytene cells in *eri- 1(mg366)*;UAD-2::GFP worms fed the indicated RNAi bacteria. **f,** Deep sequencing of total small RNAs of the indicated adult animals. Blue dashed lines indicate piRNAs. **g,** Scatter plots depicting the expression levels of Cluster Ⅰ, Cluster Ⅱ, and out-Cluster piRNAs in the wild-type (x-axis) and *tofu-3* mutant strains (y-axis), normalized as reads per million+1. **h,** Expression levels of total piRNAs from LG IV of the indicated adult animals cultured at 20°C, normalized as reads per million+1. **i-k,** Subcellular localization of UAD-2::GFP and mCherry::H2B in pachytene cells from WT, *tofu- 3(ust235)* and t*ofu-3(tm3068)* mutant worms.

**Extended Data Fig. 7.**
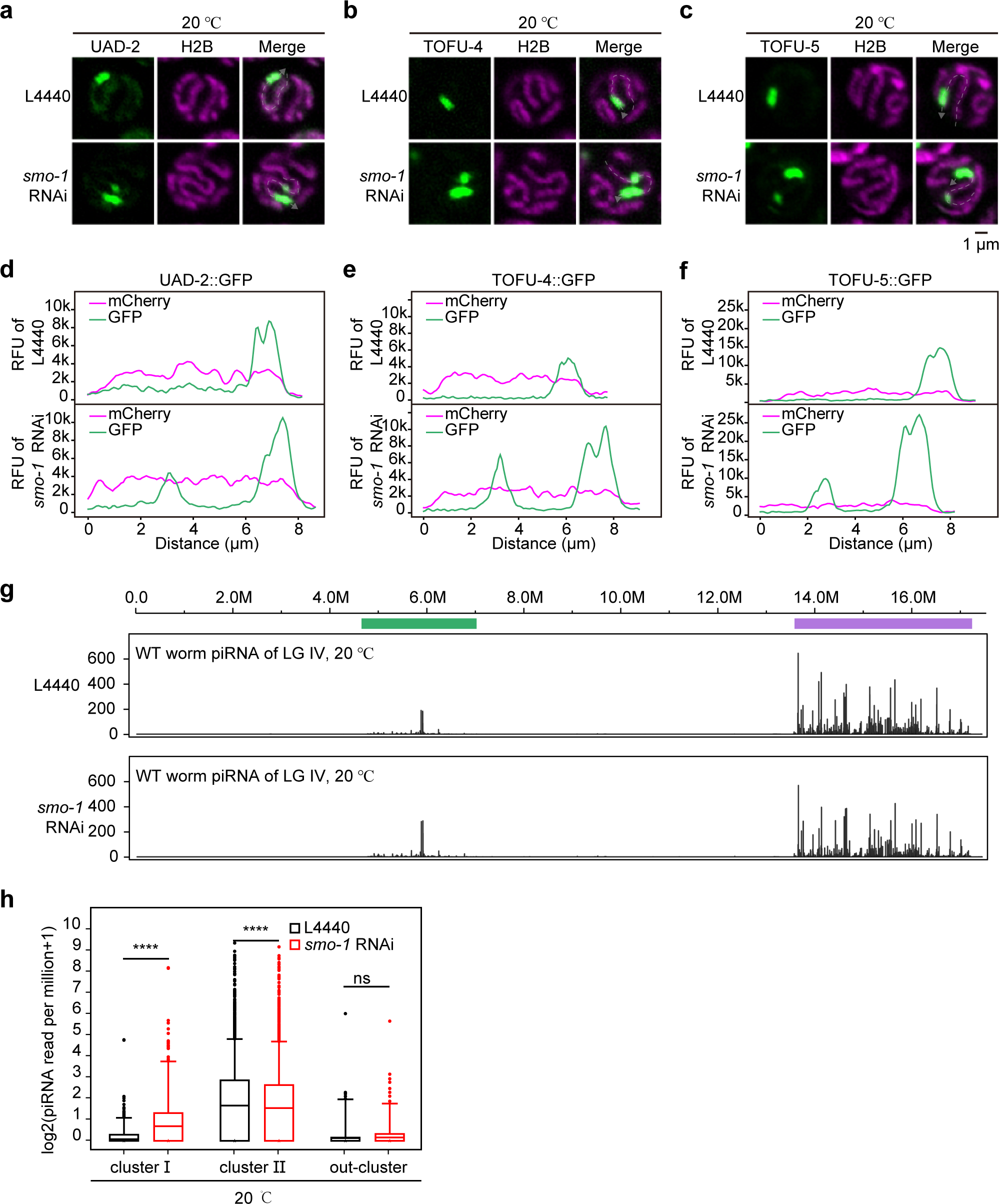
SMO-1 suppresses piRNA expression. a-c, Subcellular localization of UAD-2::GFP, TOFU-4::GFP and TOFU-5::GFP with mCherry::H2B in pachytene cells of WT worms fed L4440 and *smo-1* dsRNA at 20°C. **d-f,** Relative fluorescence unit intensity (RFU) indicated by dashed lines along chromosome Ⅳ of (a-c). **g,** Expression levels of total piRNAs from LG IV in specified adult animals grown at 20°C, normalized as reads per million+1. **h,** Boxplots illustrating log_2_(piRNA reads per million+1) for the indicated adult animals. Statistical significance was determined using a paired-sample t test.

## Notes

### Competing Interest Statement

The authors have declared no competing interest.

